# Exploring the Role of Spatial Confinement in Immune Cell Recruitment and Regeneration of Skin Wounds

**DOI:** 10.1101/2023.04.30.538879

**Authors:** Yining Liu, Alejandra Suarez-Arnedo, Eleanor Caston, Lindsay Riley, Michelle Schneider, Tatiana Segura

## Abstract

Microporous annealed particle (MAP) scaffolds are injectable granular materials comprised of micron sized hydrogel particles (microgels). The diameter of these microgels directly determines the size of the interconnected void space between particles where infiltrating or encapsulated cells reside. This tunable porosity allows us to use MAP scaffolds to study the impact of spatial confinement (SC) on both cellular behaviors and the host response to biomaterials. Despite previous studies showing that pore size and SC influence cellular phenotypes, including mitigating the macrophage inflammatory response, there is still a gap in knowledge regarding how SC within a biomaterial modulates immune cell recruitment *in vivo* in wounds and implants. Thus, we studied the immune cell profile within confined and unconfined biomaterials using small (40 μm), medium (70 μm), and large (130 μm) diameter spherical microgels, respectively. We discovered that MAP scaffolds imparted regenerative wound healing with an IgG1-biased Th2 response. MAP scaffolds generated from 130 μm diameter microgels have a median pore size that can accommodate ∼40 µm diameter spheres induced a more balanced pro-regenerative macrophage response and better wound healing outcomes with more mature collagen regeneration and reduced levels of inflammation.

## Main Text

When designing biomaterials for clinical applications, the performance of these platforms hinges upon their interaction with the host immune system^1–3^. A failure to engage and incorporate the correct immune response can lead to a foreign body response and the subsequent rejection of the materials. This is due to a cascade of cellular responses^4^, including persistent inflammation, a build-up of foreign body giant cells, and fibrosis at the implant site. To improve the biocompatibility of biomaterials and avoid undesired immune reactions, a wide range of design parameters were researched, such as surface modifications (e.g., a change in hydrophobicity or surface charge to reduce protein or cell absorption^5, 6^, incorporating cell adhesive ligands to selectively engage immune cells^7, 8^) and stiffness^9,10^. Specifically, altering material porosity by varying the size of the void space inside biomaterial scaffolds has shown promise in mitigating immune response and promoting better tissue integration^11–13^. Having an interconnected network of void space inside the scaffold allows for easy traversal of cells and diffusion of biological factors^14^, and it leaves room for the development of vasculature and stroma^15–17^. The size and shape of the void space also apply spatial constraints onto the infiltrating cells, thereby modulating their behavior and phenotypes^18^.

Only a handful of studies examined the role of material porosity and spatial confinement (SC) on cellular behaviors and tissue regeneration *in vivo*. For example, researchers observed the most blood vessel formation with minimal fibrotic response in poly (2-hydroxyethyl methacrylate-co-methacrylic acid) (pHEMA) scaffolds with pore diameters of 30 or 40 μm following 2-week cardiac implantation in rats, compared to scaffolds with 20 μm pores or no pores^12^. The existence of porous architectures in these scaffolds increased the macrophage mannose receptor (MMR, pro-regenerative marker) level in nitric oxide synthase 2 (NOS2, pro-inflammatory marker) expressing macrophages^12^. A similar study with the same pHEMA system uncovered that at 3 weeks post-implantation in mice, the pro-regenerative macrophage marker expression (MMR and scavenger receptor B I/II) was lower (<50%) inside 34 μm pores than in 160 μm pores^13^. Another work explored the influence of pore size-mediated macrophage polarization on angiogenesis^15^. They reported that in a subcutaneous implantation model, collagen scaffolds with average pore sizes of 360 μm induced more vascularization and recruited more VEGF+ cells and fewer pro-inflammatory macrophages, compared to those with 160 μm pores. These findings contributed to the understanding that SC within scaffolds of different pore sizes can guide cellular infiltration and mitigate immune cell responses. Nevertheless, the results did not consistently associate one particular optimal pore size with a preferable immune response, nor did they cover a time course of the dynamic process of the material-cell interactions.

Granular scaffolds are an emerging platform for studying how SC impacts cellular behaviors and the host response to biomaterials^19–22^. These materials are composed of microgel building blocks, which can be individually designed and collectively form constructs with desired mechanical and biochemical properties^19, 23^. The bottom-up design of granular materials offers many appealing characteristics, particularly injectable porosity. The particle size and subsequent pore size between particles are highly tunable, thereby enabling users to engineer SC within granular scaffolds^24^. In recent years, granular scaffolds have been leveraged in several applications as *in vitro* cell culture models as well as *in vivo* acellular and cell/molecule delivery therapeutics^25–27^. As this emerging platform gets widely utilized, understanding how SC within granular scaffolds modulates immune cell recruitment *in vivo* also empowers future designs to optimize particle size and induce specific cellular responses for target applications.

Microporous annealed particle (MAP) scaffolds are a type of granular material comprised of interlinked micron sized hydrogel particles (microgels) resulting in a void space network surrounding the microgels^28^. Initially designed to modulate the host response and improve cellular infiltration in wound healing^29^, MAP scaffolds soon demonstrated potential in many applications ^21, 30–41^. The modular nature of MAP scaffolds offers enormous tunability in not only the individual microgel design but also the homogenous or heterogenous microgel assembly into the bulk scaffold^32^. When engineering MAP scaffolds, the size of the microgels composing MAP scaffolds dictates the internal landscape of the void space and the resulting SC that is sensed by cells^20, 42, 43^ In this study, we explored the *in vivo* cellular response to MAP scaffolds with various particle sizes at different time points in a subcutaneous implant model and a wound healing model. We discovered that MAP scaffolds with smaller pore sizes hindered the initial cell infiltration and required more cell-guided degradation, which led to constructive innate and adaptive immune responses. We employed a multifaceted flow cytometry phenotyping approach to quantify the infiltrating cell types and phenotypes in MAP scaffolds in both models. We concluded that the pore size-dependent recruitment and phenotype modulation of macrophages and other immune cells impacted the pro-healing effect observed in MAP scaffolds. Scaffolds comprised of large microgels with unconfined pore sizes resulted in a less inflammatory and more pro-regenerative responses.

### 40 μm, 70 μm, and 130 μm MAP scaffolds were chemically and mechanically identical

Spherical microgels of different sizes were generated using different microfluidic devices with various run speeds (Figure 1a). The diameter of small microgels was around 40 μm, medium around 70 μm, and large 130 μm ^44^. An analytical software (LOVAMAP) was developed in our lab to study and characterize different aspects of the void space inside MAP scaffolds for optimizing scaffold designs^42^. This software segments the void space into natural pockets of open space (3D-pores) that are delineated at ‘doors,’ which are the narrower regions of void space between touching or nearby particles (Figure 1b). When the doors are on the surface of the scaffold, they are referred to as “entry doors” that infiltrating cells may use to enter the scaffold. We observed through confocal images that MAP scaffolds formed by different-sized microgels (abbreviated as 40 μm, 70 μm, 130 μm MAP scaffolds) had consistent void fraction (25-35%) but very distinct internal structures (Figure 1c)^44^. Using LOVAMAP to further characterize the interior of MAP scaffolds, we observed that the diameter range of the largest enclosed sphere within each 3D-pore in 40 μm MAP scaffolds mostly fell below the average size of a myeloid cell (Figure 1d). The same trend could also be observed for the diameter range of entry doors on the surface of 40 μm MAP scaffolds, whereas the diameters of most entry doors in 70 μm and 130 μm MAP scaffolds were larger than the diameter of a myeloid cell (Figure 1e). The microgel size-dependent shift in the pore and entry door profiles indicated that when these MAP scaffolds are used *in vivo*, they apply different degrees of spatial confinement on the infiltrating cells. We matched material stiffness for all three scaffold types so that the difference in cellular response could only be attributed to the change in spatial confinement^44^.

**Figure 1:**
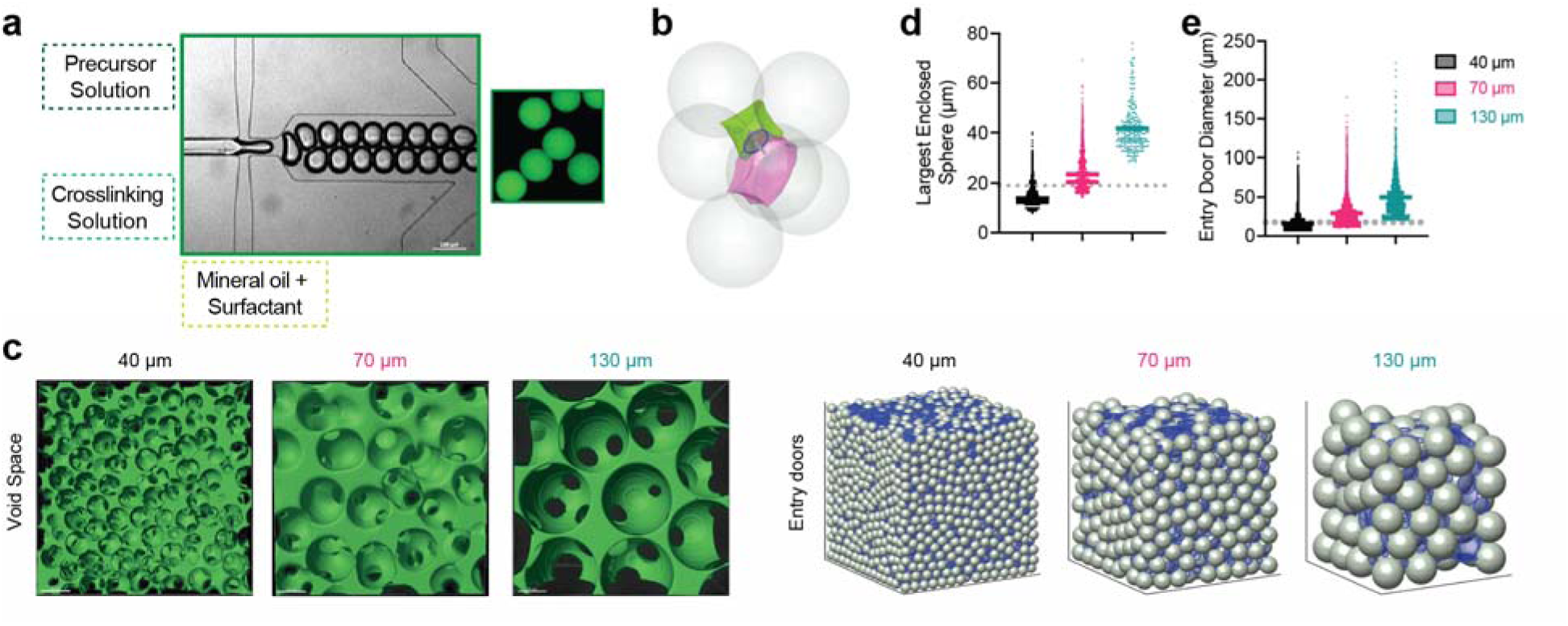
40 μm, 70 μm, and 130 μm MAP scaffolds were chemically and mechanically identical. a, scheme illustrating microgel formation using a microfluidic water-in-oil emulsion system. A precursor solution and a crosslinker solution are fused and segmented into droplets, which are then crosslinked into microgels via Michael addition. b, sample green and pink 3D-pores separated by a door (blue circle) and surrounded by six microgels. c, (Left) fluorescent images showing void space; (Right) entry doors (blue circles) for 40 μm, 70 μm, and 130 μm simulated MAP scaffolds. d, the diameter of the largest enclosed sphere within each 3D-pore for simulated 40 μm, 70 μm, and 130 μm MAP scaffolds compared to the diameter of bone marrow-derived macrophages (BMDM) in 3D culture (grey dotted line). e, the diameters of all entry doors for simulated 40 μm, 70 μm, and 130 μm MAP scaffolds compared to the diameter of BMDM in 3D culture (grey dotted line). Statistical analysis: one-way ANOVA.

### Immune cell recruitment and response followed a size-dependent manner in the subcutaneous implantation model

To explore the role of spatial confinement on immune cell recruitment in a non-traumatic setting, we characterized the infiltrating immune cell profile in a subcutaneous implantation model. Each mouse received three injections of 50 µL hydrogel (one of each type, 40 μm, 70 μm, and 130 μm MAP scaffolds) on the back. After 1-, 4-, 7-, 14- and 21-days post-implantation, the infiltrated cells were extracted, and their phenotypes were analyzed by an 11-color innate immune cell panel (Figure 2a). This time window was chosen to explore the full spectrum of the foreign body response (FBR): days 1 and 4 represent the initial acute inflammatory phase and are followed by a late inflammatory phase around 7 to 14 days; the resolution phase occurs after 21 days. The total infiltrated live cell number and CD45+ cell number were similar among all MAP scaffolds groups (Figure 2b), which indicated that a smaller entry door size didn’t pose a significant barrier to the cell infiltration. Macrophages and FcεRI+ cells (such as Langerhan cells, monocytes, mast cells, and other dendritic cells) dominated the immune cell infiltration^45^ (Figure 2c). The immune cell recruitment followed a size-dependent manner. 130 μm MAP scaffolds recruited significantly more neutrophils within one day while 40 μm MAP scaffolds had elevated FcεRI+ cell percentage and 70 μm MAP scaffolds had a higher ratio of basophils (Figure 2d). Preceded by an elevation in infiltrating monocytes at day 4, the macrophage percentage peaked at day 7 in all the groups and most significantly in 130 μm MAP scaffolds (Figure 2d). Eosinophil level was increased at day 7 and more eosinophils were accumulated in 40 μm MAP scaffolds than 130 μm MAP scaffolds. As the FBR moved toward resolution, the percentage of FcεRI+ cells increased over time in all scaffolds and again was the highest in 130 μm MAP scaffolds (Figure 2d). The general immune response towards all MAP scaffolds was active but constructive, with an improved level of collagen deposition, granulation tissue, and vascularization 21-day post-implantation (Supplementary Figure 1a).

**Figure 2.**
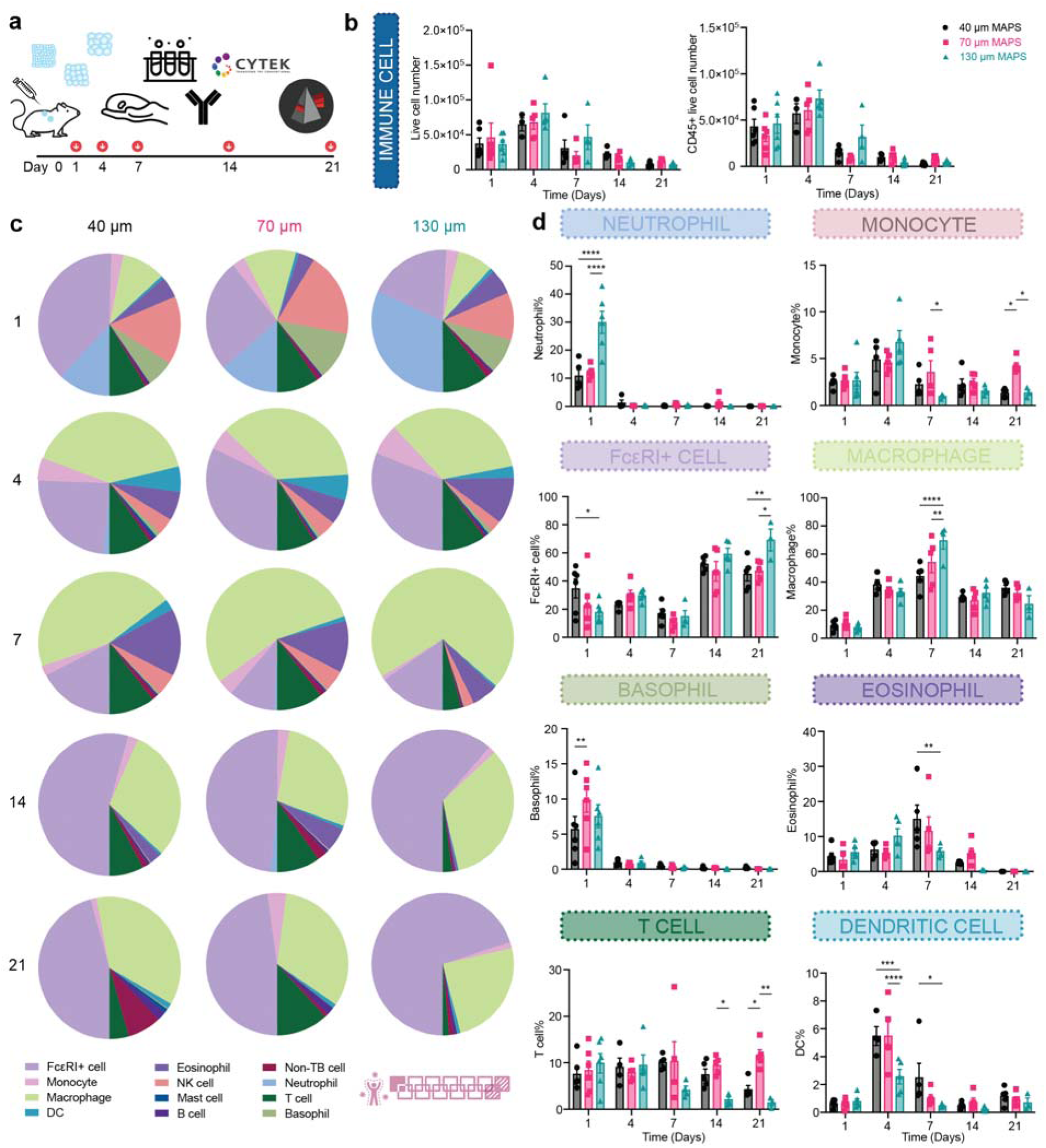
Immune cell recruitment and response followed a size-dependent manner in the subcutaneous implantation model. a, Scheme illustrating the experiment timeline. After the initial injections, implant extraction and flow cytometry were performed at designated time points (days 1, 4, 7, 14, 21). b, the total number of live cells (Zombie NIR-) and CD45+ immune cells (CD45+). c, pie charts of myeloid cell abundancy across 5 time points for 40 μm, 70 μm, and 130 μm MAP scaffolds. Each number was an average of the results from 5 mice. d, neutrophil, monocyte, FcεRI+ cell, macrophage, Basophil, eosinophil, T cell and dendritic cell percentages among all CD45+ live cells. Statistical analysis: two-way ANOVA with Šídák’s multiple comparisons test made between 40 μm, 70 μm, and 130 μm MAP scaffolds groups only when there was a significance in the interaction term of scaffold type x time. * p<0.05, ** p<0.01, *** p<0.001, **** p<0.0001. Error bars, mean ± s.e.m., n = 6 mice per group for day 1 and n = 5 mice per group for the other time points. The pink symbol in the middle bottom of the graph stands for the 13-color innate cell panel used in this figure.

### 130μm MAP scaffolds induced mature collagen regeneration and reduced inflammation in skin wounds

To see how the size-dependent immune cell recruitment in MAP scaffolds would impact wound healing outcomes, we compared the effect of MAP scaffold treatment against a clinically used standard treatment Woun’Dres in a full-thickness excisional wound healing model in mice. To characterize the quality of tissue regeneration, we assessed the collagen architecture and maturity with Picro-Sirius red (PSR) staining and Masson’s Trichrome (MT) staining (Figure 3a, Supplementary Figure 2a, Supplementary Figure 3a). In PSR, the amount of collagen and its architecture can be visualized and characterized by bright field and polarized light^46^. In MT, the mature collagen appears in deep blue and has a basket weave-like network whereas newly regenerated collagen fibers are light blue in color and have more aligned fiber orientation^47, 48^. 130 μm MAP scaffolds treated wounds appeared to have a structure more closely resembling normal skin around the wound bed, showing better collagen regeneration with a higher collagen percentage and longer average fiber length than Woun’Dres group (Figure 3a-c). The 130 μm MAP scaffold group had the highest average epidermis to dermis (E/D) ratio, which was statistically greater than the rest of the groups and closer to the normal skin baseline (Figure 3e). Both 40 μm and 130 μm MAP scaffold groups had a lower afollicular percentage (greatest distance between two follicular structures in the wound bed divided by the total length of the wound) than Woun’Dres group (Figure 3f), indicating a smaller “true scar” region within the wounds.

**Figure 3:**
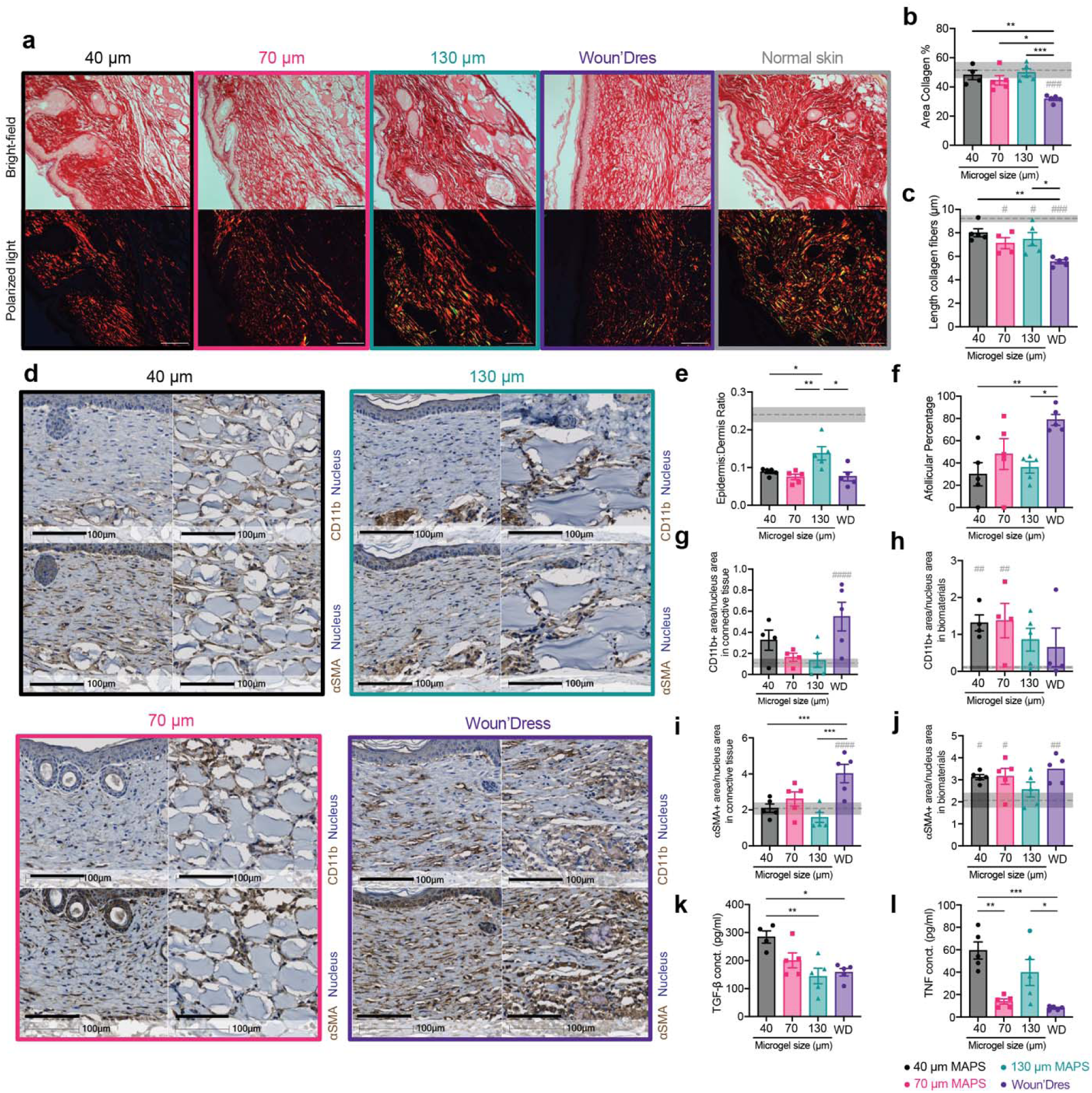
130 μm MAP scaffolds induced mature collagen regeneration and reduced inflammation level in the skin wound. a, representative pictures of 21-day skin wound samples with Picro-Sirius Red staining (top row, bright field image, scale bar, 100 μm; bottom row, image with polarized light, scale bar, 100 μm). b, percentage of collagen (bright field area) in the regions of interest. c, average length of collagen fibers in the regions of interest. d, representative pictures of 21-day skin wound samples with immunohistochemical staining of CD11b and α-SMA (left: connective tissue in the wound bed, scale bar, 100 μm; right: remaining biomaterials, scale bar, 100 μm). e, the ratio of epidermis thickness to dermis thickness. The dotted line and the grey area stand for the average number and the range for normal skin. f, afollicular percentage in the wound bed. g-h, the percentage of CD11b+ area as a ratio of the nucleus area in the connective tissue and in the remaining biomaterials. i-j, the percentage of α-SMA + area as a ratio of the nucleus area in the connective tissue and in the remaining biomaterials. k-l, ELISA results of TGF-β and TNF concentrations inside the hydrogel implants 21 days post-wounding. Statistical analysis: two-way ANOVA with Šídák’s multiple comparisons test made between treatment groups only when there was a significance in the interaction term of treatment type x time. Dunnet method was used to compare each treatment against normal skin baseline (gray pond). ∗/# p < 0.05, ∗∗/## p < 0.001, ∗∗∗ p < 0.001, ∗∗∗∗/#### p < 0.0001. Error bars, mean ± s.e.m., n = 5 mice per group with some data points removed due to experimental reasons.

Improved regeneration of the skin was also supported by the timely resolution of inflammation and matrix remodeling. CD11b+ immune cell accumulation, an indication of the general inflammation level, in the connective tissue of MAP scaffolds-treated wounds was close to that of normal skin, whereas the Woun’Dres group still had a significantly higher level than the baseline (Figure 3d, g). In the remaining biomaterials in the wound bed, 130 μm MAP scaffolds and Woun’Dres treatment both had a CD11b+ cell amount not significantly different than the basal level (Figure 3d, h). A similar trend was also confirmed by the histological assessment that the general inflammation level on day 21 was lower in 130 μm MAP scaffolds when compared to Woun’Dres groups (Supplementary Figure 2b). Myofibroblasts, a major cell type secreting extracellular matrix (ECM), are responsible for depositing and replacing collagen during wound healing and fibrosis^49^. An excess of myofibroblasts, characterized by the expression of alpha-smooth muscle actin (α-SMA), at the resolution phase of wound healing is associated with fibrosis^50^. In the newly formed connective tissue, MAP scaffolds groups restored the α-SMA+ myofibroblast level to the baseline while Woun’Dres group remained at a significantly higher level of α-SMA+ cell accumulation (Figure 3d, i). Within the remaining biomaterials, only the 130 μm MAP scaffolds group had a α-SMA+ cell level close to that in normal skin (Figure 3d, j), suggesting a better resolution of matrix remodeling.

ELISA results reviewed a size-specific cytokine profile at day 21. Notably, TGF-β, a cytokine associated with tissue regeneration and fibrosis^51, 52^, was significantly increased in 40 μm MAP scaffolds treated wounds compared to 130 μm MAP scaffolds treated wounds, which pointed to a resolution of remodeling in 130 μm MAP scaffolds group after 21 days (Figure 3k). TNF, a potent inflammatory cytokine, was elevated significantly both in 40 μm and 130 μm MAP scaffold groups (Figure 3l). IL-1β, a macrophage activation cytokine, was generally expressed in all the treatment groups (Supplementary Figure 2m). The key cytokine for Th2 immune response IL-4 was only expressed at a relatively low level in all groups (Supplementary Figure 2n). Collectively, these results indicated a refined skin regeneration was promoted in wounds treated by a less confined scaffolds.

### Immune cell recruitment followed a size-dependent manner in skin wounds

An effective and efficient wound repair requires the coordination of many different cell types, especially the immune cells during the inflammation phase. Given the improved wound healing outcomes we observed in MAP scaffold-treated wounds, especially with 130 μm MAP scaffolds, we further explored the immune cell profile in the skin wounds. We sampled a 5 mm area around the initial wound site at days 1, 7, and 21. This time window was chosen to explore the acute immune response towards the materials (days 1 and 7) and the resolution phase 21-day post-wounding (Figure 4a). The total infiltrated live cell number and CD45+ cell number were similar among all groups, again confirming that the size range of the entry door in MAP scaffolds was still large enough to allow sufficient cell infiltration (Figure 4b). To get an overview of the cellular infiltration, a representative profile in all treatment groups was mapped out (Figure 4c). Neutrophils and monocytes dominated the immune cell infiltration at day 1. 40 μm and 130 μm MAP scaffolds recruited more neutrophils than 70 μm MAP scaffolds whereas 70 μm MAP scaffolds attracted the highest percentages of monocytes, macrophages, and FcεRI+ cells (Figure 4d), which are the main immune modulators and antigen presenting cells (APCs) in the skin wound. At day 7, neutrophils and basophils took up most of the immune cells while T cells and basophils were the major cell types in the immune infiltrate 21-day post-wounding (Figure 4c). Since we observed size-dependent immune cell recruitment (Figure 4d), specifically at day 1, we sought to explore further the impact of spatial confinement on immune cell phenotype and focus on macrophages, a key immune modulator during wound healing.

**Figure 4:**
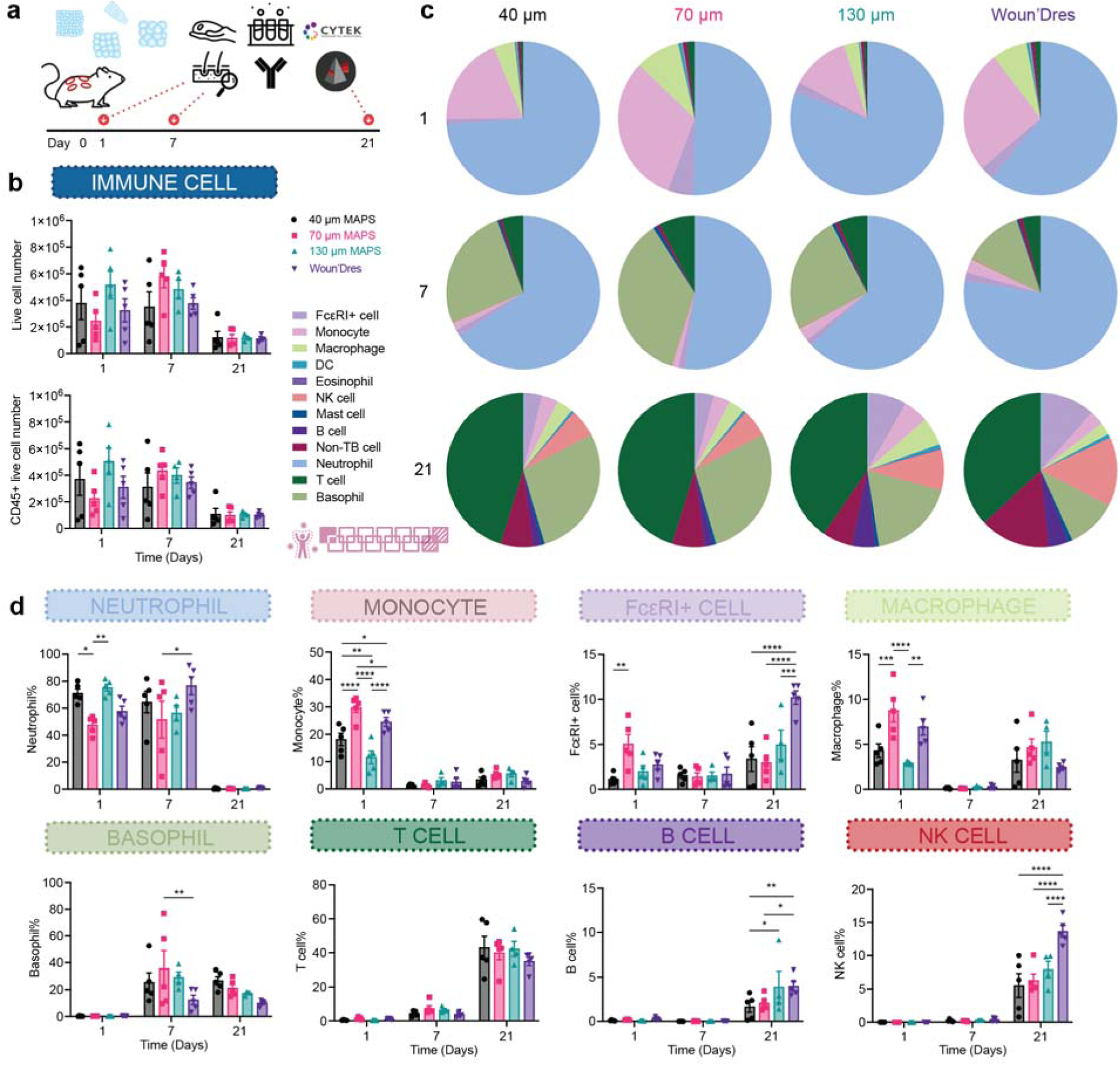
Immune cell recruitment and response followed a size-dependent manner in the wound healing model. a, Scheme illustrating the experiment timeline. After the initial wounding and biomaterial treatment, skin extraction and flow cytometry were performed at designated time points (days 1, 7, 21). b, the total number of live cells (Zombie NIR-) and CD45+ immune cells across three time points. c, pie charts of myeloid cell abundancy across 3 time points for 40 μm, 70 μm, and 130 μm MAP scaffolds. Each number was an average of the results from 5 mice. d, neutrophil, monocyte, FcεRI+ cell, macrophage, basophil, T cell, B cell, NK cell percentages among all CD45+ live cells. Statistical analysis: two-way ANOVA with Šídák’s multiple comparisons test made between 40 μm, 70 μm, and 130 μm MAP scaffolds and wound dressing groups only when there was a significance in the interaction term of scaffold type x time. * p<0.05, ** p<0.01, *** p<0.001, **** p<0.0001. Error bars, mean ± s.e.m., n = 5 mice per group. The pink symbol in the middle right of the graph stands for the 13-color innate cell panel used in this figure.

### 130 **μ**m MAP scaffolds modulated a timely transition in pro-regenerative macrophage phenotypes

Spatial confinement of macrophages, an essential cell type in wound healing, has been shown to result in a less inflammatory phenotype^18^. We explored the change in macrophage phenotype during the wound healing process and in response to the change in spatial confinement. A significantly higher number of macrophages infiltrated 40 μm MAP scaffolds and Woun’Dres groups than 70 μm and 130 μm MAP scaffolds groups at day 1 (Figure 5a). Around 60% of these macrophages expressed Ly6C, which indicated an origin from circulating monocytes in blood (Figure 5b). The Ly6C+ macrophage percentage dropped in MAP scaffolds groups at day 7 as the cells differentiated into macrophages and assumed specific functions (Figure 5b). Almost all the macrophages were fully mature and lost Ly6C expression after 21 days (Figure 5b).

**Figure 5:**
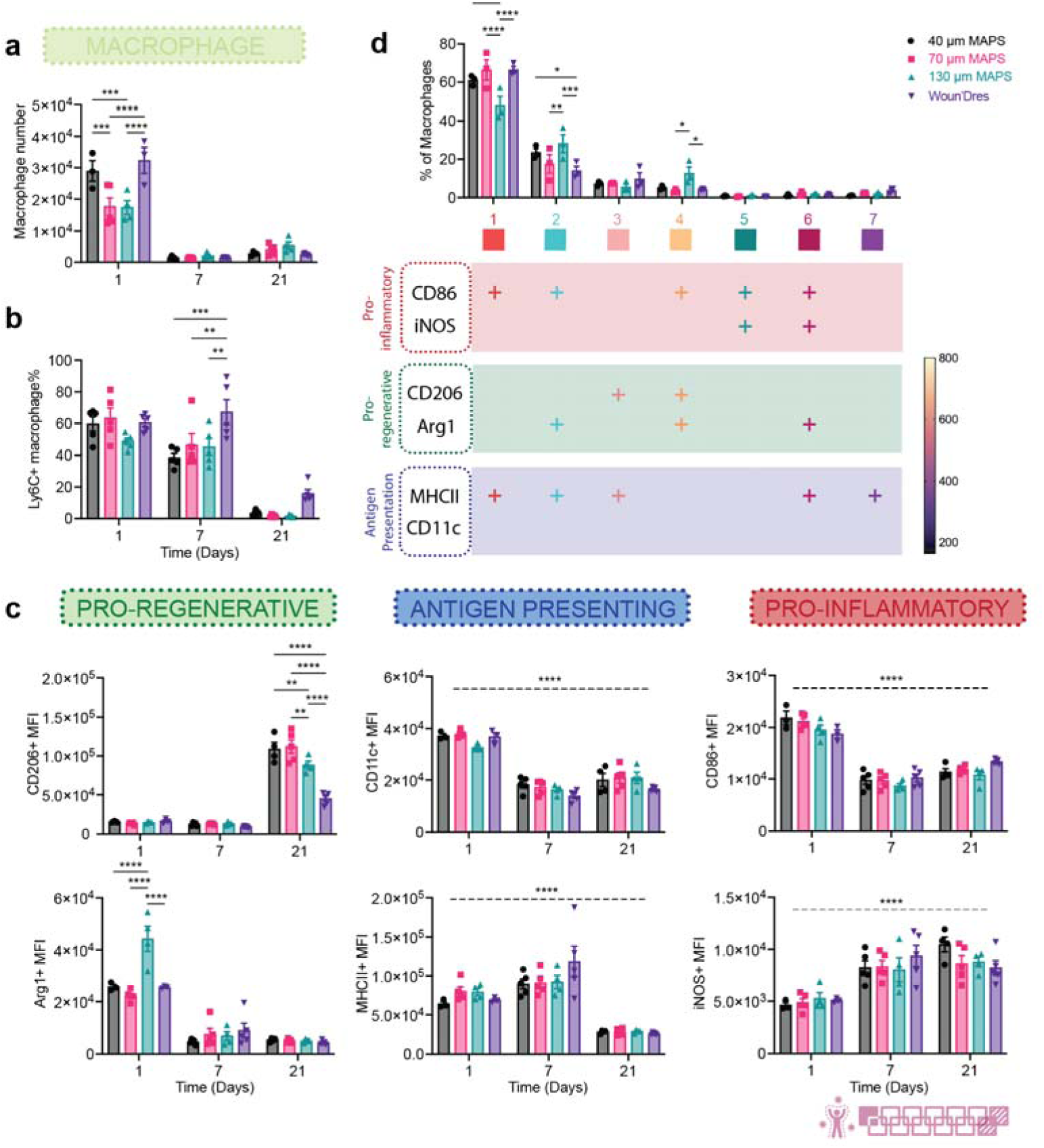
130 μm MAP scaffolds modulated a timely transition in pro-regenerative macrophage phenotypes. a, the number of macrophages in total live cells across 3 time points. b, Ly6C+ macrophage percentage in total macrophages. c, MFI of CD206, Arg1, CD11c, MHCII, CD86, and iNOS in total macrophage population over time. d, tSNE clustering of macrophages in MAP scaffolds-treated wounds on day 1. The bar graph showed the percentages each sub-population took up in the total macrophage population. The table beneath it showed the expression levels of each marker in 7 different macrophage populations. “+” indicates that more than 50% of the population expressed that marker. Statistical analysis: two-way ANOVA with Šídák’s multiple comparisons test made between 40 μm, 70 μm, and 130 μm MAP scaffolds and wound dressing groups only when there was a significance in the interaction term of scaffold type x time. * p<0.05, ** p<0.01, *** p<0.001, **** p<0.0001. Asterisks with solid line stand for comparisons between groups. Asterisks with dash line stand for significance in time. Error bars, mean ± s.e.m., n = 5 mice per group. The pink symbol in the bottom right corner of the graph stands for the 13-color innate panel used in this figure.

Macrophages in all the wounds showed a time-dependent phenotype transition (Figure 5c). Higher levels of pro-inflammatory/co-stimulatory marker CD86 and antigen-presenting marker CD11c were observed at day 1. Antigen-presenting marker MHCII was also highly expressed at days 1 and 7. The expression of these markers suggested the active antigen presentation role macrophages play during the early wound healing response. Nitric oxide (NO) release has been previously reported to significantly reduce collagen encapsulation and chronic inflammation in foreign body response^53^. The pro-inflammatory marker inducible nitric oxide synthase (iNOS), a key enzyme driving the production of immunomodulating molecule NO, remained highly expressed at days 7 and 21 as the wound moved towards resolution. Notably, both pro-regenerative markers Arginase 1 (Arg1) and CD206 demonstrated size-dependent shifts in expression. The induction of arginase activity in the wound could cause the depletion of arginine, which provides substrates indirectly contributing to collagen deposition^54^. Arg1 was highly expressed in wound macrophages at day 1, especially in the 130 μm MAP scaffolds group. CD206 levels were also elevated in all MAP scaffolds groups, 40 μm and 70 μm MAP scaffolds in particular, at day 21. A significantly lower level of CD206 as day 21 in 130 μm MAP scaffolds wounds compared to 40 μm and 70 μm MAP scaffolds groups, together with a generally tamed expression of the other five functional markers. This pointed to a resolution of macrophage response in 130 μm MAP scaffolds that might in part contribute to the better wound resolution and more mature collagen regeneration.

To further characterize the macrophage phenotype profile at day 1 (the initial acute immune response with the most macrophage accumulation), we used FlowSOM, an unsupervised algorithm based on Self-Organizing Maps, to reveal how all six functional markers were behaving on macrophages and distinguish distinct subpopulations^55^. These macrophage populations were differentiated based on the expression levels of each marker (shown in the table below the histogram plot, “+” meant over 50% population expressed that marker). The FlowSOM analysis showed that MAP scaffolds attracted similar profiles of macrophages and polarized them into a broad range of phenotypes. Notably, the pro-inflammatory markers CD86 and/or iNOS in some cases (population 2, 4, 6) were co-expressed with pro-regenerative markers CD206 and/or Arginase 1. This finding supported that macrophage phenotypes exist on a spectrum and their functions cannot be fully captured by one or two markers. A total of 7 distinct macrophage populations were identified. Population 1, which expressed CD86 and MHCII, was an antigen-presenting phenotype and the major macrophage subpopulation in all wounds. 130 μm MAP scaffolds-treated wounds recruited higher percentages of population 2 (Arg1+CD86+MHCII+, M2-biased antigen-presenting phenotype) and 4 (Arg1+CD86+CD206+, M2-biased hybrid phenotype), both expressing a high level of Arg1 and exhibiting a pro-regenerative polarization. Collectively, these observations confirmed that MAP scaffolds induced a complex macrophage response with unique expression profiles. 130 μm MAP scaffolds promoted an early pro-regenerative macrophage profile and modulated a timely resolution in macrophage response towards the end of the wound healing process.

### A pro-reparative IgG1-biased Th2 response was observed in 70 **μ**m MAP scaffolds

Since a significant amount of adaptive immune cells, especially T cells, were recruited to the wounds after material treatment, we examined whether the application of MAP scaffolds would lead to any systemic adaptive immune response. Remarkably, by day 21, all mice with MAP scaffolds treatment had developed MMP linker-specific IgG response, with 70 μm MAP scaffolds having the most drastic increase compared to the untreated control (Figure 6a). This MAP scaffolds-induced response was IgG1-biased and therefore a pro-reparative Th2 response (Figure 6b). We also investigated the adaptive immune cell profiles of draining lymph nodes (dLN) and the spleen in MAP scaffolds-treated mice 1-day, 7-day, and 21-day post-wounding. MAP scaffolds treatment also modulated T cell profile and an increase in Th2 marker GATA3 expression in dLN at day 7 and in the spleen at days 7 and 21 was observed (Figure 6c,d). Compared to the baseline, the Th1 marker Tbet level was significantly reduced in dLN at days 7 and 21 and in almost all groups except 70 μm MAP scaffolds at day 7 in the spleen (Figure 6c,d). The evidence provided here demonstrates that a pro-regenerative Th2 adaptive immune response can be engaged by synthetic pro-regenerative scaffolds and its magnitude was modulated by the size of the microgels^26^ (Figure 6c,d, Supplementary Figure 3).

**Figure 6:**
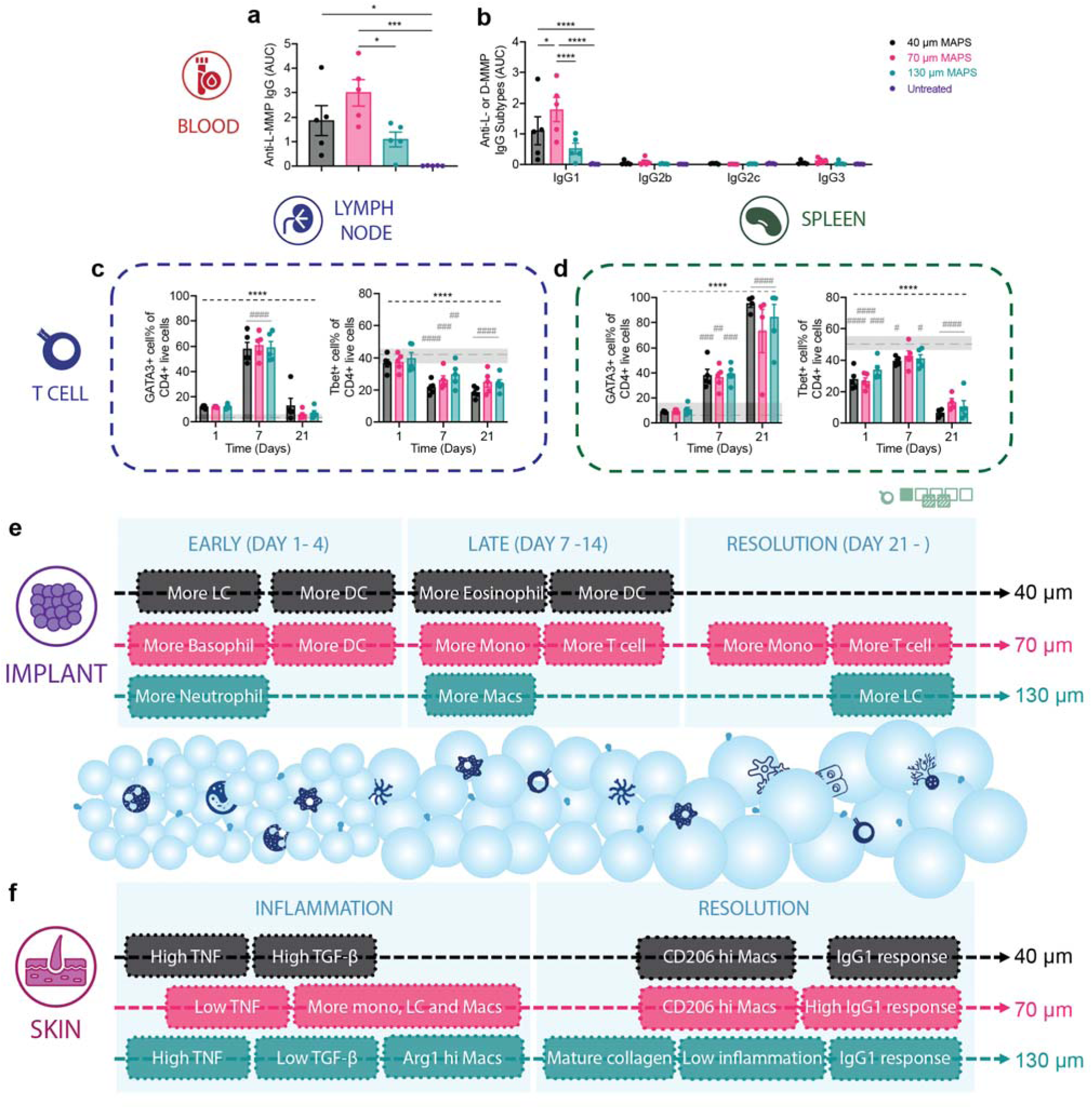
A pro-reparative IgG1-biased Th2 response was observed in 70 μm MAP scaffolds. a and b, ELISA results of total anti-L-MMP (peptide crosslinker) IgG level and the anti-L-MMP IgG subtypes in mice treated with 40 μm, 70 μm, and 130 μm MAP scaffolds after 21 days. c and d, T cell profiles in draining lymph nodes and spleen of mice treated with 40 μm, 70 μm, and 130 μm MAP scaffolds across 3 time points. e and f, schematic illustrations of the immune responses to 40 μm, 70 μm, and 130 μm MAP scaffolds in the subcutaneous implantation model and the wound healing model. Statistical analysis: two-way ANOVA with Šídák’s multiple comparisons test made between 40 μm, 70 μm, and 130 μm MAP scaffolds groups only when there was a significance in the interaction term of scaffold type x time. After a two-way ANOVA, Dunnet method was used to compare the experiment groups with the baseline control group. * p<0.05, ** p<0.01, *** p<0.001, **** p<0.0001. Asterisks with solid line stand for comparisons between MAP scaffolds. Asterisks with dash line stand for significance in time. Error bars, mean ± s.e.m., n = 5 mice per group but with some data points removed due to experimental reasons. The green symbol beneath panel d stands for the 7-color T cell panel used in panel c, d of this figure.

## Discussion

Pore size has long been a key design parameter for biomaterials. As the entry points into scaffolds, the pore size on the surface of materials poses limitations for infiltrating cells and ingrowing tissues. A small pore size, usually on the nanoscale or below 10 microns, is known to prevent cellular infiltration and is more prone to induce a foreign body response^13, 17^. On the other hand, larger pore sizes of tens to hundreds of microns could be beneficial for nutrient transport and the growth of connective tissue and blood vessels^13, 15, 56^, which supports implant success rate in the long run. Once the infiltrating cells are in the interior of the scaffolds, the size and architecture of the void space influences cell morphology, phenotype, and activity by imposing spatial constraint and confinement. Spatially confining immune cells, such as macrophages, can modulate their phenotype and their inflammatory response^18^. Scaffolds with an optimized pore size should support the initial influx of immune cells and offer sufficient spatial confinement to modulate cell phenotype while accommodating structural regeneration.

In this study, we demonstrated that microporous annealed particle scaffolds (MAP scaffolds) consisting of different-sized microgels (40 μm, 70 μm, and 130 μm diameter) imparted regenerative wound healing with an IgG1-biased Th2 response when compared to clinical standard treatment. 130 μm MAP scaffolds with a pore size that can accommodate ∼40 µm diameter spheres induced mature collagen regeneration and reduced levels of inflammation in the skin wound. Pore size-dependent recruitment and phenotype modulation of macrophages and other immune cells impacted the pro-healing effect observed in MAP scaffolds. These findings suggest that pore size can be specifically engineered to modulate the functions of infiltrating cells and trigger regeneration rather than fibrosis. Future directions combining the MAP scaffolds platform with other immune or pro-regenerative factors can further leverage its potent immunomodulatory potential for optimal skin regeneration.

## Author contributions

YL made substantial contributions to the conceptualization and design of the work, acquisition, analysis, and interpretation of data, as well as writing the manuscript. ASA participated in the acquisition, analysis, and interpretation of data for both the subcutaneous and wound healing models, as well as drafting the manuscript. EC sectioned, stained, and performed histological quantification of wound samples found in figure 3 and supplementary figure 2. LR computationally analyzed the scaffolds and provided analysis, interpretation, and writing. MS oversaw and performed histology assessment that contributed substantially to figure 3 and supplementary figure 2. All the authors discussed the results and contributed to writing portions of the manuscript and editing the manuscript. TS conceptualized and conceived the project, provided guidance and discussion throughout, and made substantial contributions to data analysis, figure preparation, and manuscript editing.

## Acknowledgements

We would like to thank the National Institutes of Health (R01AI152568) and all the members of the Segura lab at Duke University for their support. A special thank you to Shamitha Shetty and Christopher Lloyd at Duke University for their generous help and delightful company. Thank you to April Espinoza for helping with part of the experiment. Thank you to our lab alumni Jingyi Xia for assistance in material characterization. Thank you to Patrick Duncker, Ph.D. at Cytek Biosciences for providing technical support on spectral flow cytometry and sharing insights on data interpretation. Thank you to Minerva Matos-Garner, MA from Duke Engineering Graduate Communications and Intercultural Programs for her constructive suggestions on the manuscript and her warm support during the writing process.

## Data availability

The data that support the findings of this study are available from the corresponding authors upon reasonable request. Source data are provided with this paper.

## Methods

### Microparticle generation and purification

Microfluidic devices and microgels were produced as previously described. Briefly, 8 arm PEG-VS microgels were formulated at a final concentration of 5 wt% (w/v) PEG-VS (JenKem technology) in 0.3 M triethanolamine (Sigma) with 500 µM RGD (Ac-RGDSPGERCG-NH2, GenScript), 500 µM K-peptide (Ac-FKGGERCG-NH2, GenScript) and 500 µM Q-peptide (Ac-NQEQVSPLGGERCG-NH2, GenScript) and crosslinked at a 0.6 VS to thiol ratio with di-thiol matrix metalloproteinase sensitive peptide (GenScript) plus 10 μM AlexaFluor-647 (ThermoFisher). For 40 μm and 70 μm microgel generation, we used a four-inlet microfluidic device. The aqueous solutions (precursor and crosslinker) did not mix until droplet segmentation on the microfluidic device. The pinching oil phase was a heavy mineral oil supplemented with 1% v/v Span-80. Downstream from the segmentation region, a second oil inlet with a high concentration of Span-80 (5% v/v) and Triethylamine (3% v/v) was added and mixed to the flowing droplet emulsion. These microgels were collected and allowed to react overnight at room temperature. For 130 μm microgel generation, we used a two-inlet microfluidic device and pre-mixed the precursor solution with an equal volume of the crosslinker solution. These microgels were collected in Span-80 (5% v/v) and Triethylamine (3% v/v) oil bath and allowed to gel overnight at room temperature to form microgels. The microgels were then purified by repeated washes with HEPES buffer (pH 8.3 containing 1% Antibiotic-Antimycotic) and centrifugation. The purified microgels were stored in HEPES buffer (pH 8.3 containing 1% Antibiotic-Antimycotic and 10 mM CaCl_2_) at 4 °C.

### Generation of MAP scaffolds from microgels and mechanical testing

The storage buffer in microgels was properly removed by centrifuging at 22 000 G for 5-20 minutes and taking off the supernatant. For every 50 µL of dried microgels, 2 µL of Factor XIII (250 U/mL) and 1 µL of Thrombin (200 U/mL in 200 mM Tris-HCl, 150 mM NaCl, 20 mM CaCl2) was added and mixed via thorough pipetting. A 30-minute incubation time at 37°C is required to form a bulk hydrogel scaffold. Rotational rheometry (Anton-Parr, MCR301) was used to measure the storage moduli of the scaffolds with a frequency sweep test (0.1-100 rad/s shear frequency and 1% strain amplitude).

### Microgel size and void volume measurement

Microgel size was calculated from at least three 4X Nikon Ti Eclipse pictures per batch of microgels and an average of at least five batches of microgels using a custom MATLAB code. For scaffold void volume measurement, three scaffolds were made for each type of scaffold (40 μm, 70 μm and 130 μm MAP scaffolds). Three regions of interest were randomly selected inside each scaffold and a 140 μm thick z-stack sample (509 z-slices, 0.275 um each step) was taken in each ROI at an objective of 40X. IMARIS (Oxford instruments) was used to analyze the images for void volume fraction calculation.

### Subcutaneous implantation

7-12-week-old male C57BL/6 mice (Jackson Laboratory) were anesthetized with 3.0% isoflurane and maintained at 1.5-2.0% isoflurane. The microgels and crosslinker solution were thoroughly mixed and loaded in a 1 cc syringe with a 29-gauge needle. Each mouse received three injections of 50 µL hydrogel (one of each, 40 μm, 70 μm, and 130 μm MAP scaffolds) on the back. After injection, mice were monitored until full recovery from the anesthesia. All procedures were approved by the Duke University Institutional Animal Care and Use Committee and followed the NIH Guide for the Care and Use of Laboratory Animals.

### Implant extraction and flow cytometry study

At designated time points, the implants were extracted and diced finely prior to enzymatic digestion with the digestion solution (200 U/ml Collagenase IV and 125U/ml DNase I) in RPMI media for 15 minutes at 37°C. The resulting material was filtered through a 70 μm cell strainer and washed once with 1xPBS to get a single-cell suspension. These cells were then stained with Zombie NIR (BioLegend) for 15 minutes at room temperature to access the viability and blocked with Fcr Blocking Reagent (Miltenyi Biotec) for 10 minutes on ice, followed by surface marker staining for 30 minutes on ice. For intracellular staining, a commercial Intracellular Fixation & Permeabilization kit (Thermo Fisher) was used following the instructions. After staining, samples were washed and resuspended in 150 ul flow buffer (1x PBS, 1 mM EDTA, 0.2% BSA, 0.025% proclin) and analyzed on the Cytek NL-3000 Flow Cytometer. Data was acquired using SpectroFlo software and analyzed using FlowJo™ v10.8 Software (BD Life Sciences). Relative abundance of immune cell populations was determined as a fraction of CD45+ viable cells (Gated first on scatter FSC x SCC, then doublet discrimination via FSC-A x FSC-H, prior to Viability Zombie NIR-/CD45+).

Clustering of flow cytometry data was completed by concatenating an equal number of cells pooled from all biological replicates into one file and clustering with the tSNE (t-distributed stochastic neighbor embedding) plugin for 1000 iterations, operating at theta = 0.5. Data are displayed as user-gated populations graphed against their respective X and Y tSNE coordinates. Downstream clustering was also performed with the FlowSOM algorithm on the same concatenated files and the main immune subsets were phenotypically isolated by choosing 5-8 metaclusters. Each subset was further identified by the expression or absence of different phenotypical markers.

### Wound healing study

Female and male SKH-1 Elite mice (10-12 weeks old) were purchased from Charles River and housed in a centralized animal facility at Duke University. The excisional splinted wound protocol represents an established methodology previously reported by others^57^ and is detailed in our previous publication^58^. Briefly, mice were anesthetized with 4% isoflurane and maintained at 1.5–2% isoflurane for the duration of the surgery. To prevent additional pain and hypothermia, mice were placed on a warming pad and Buprenorphine SR-Lab (ZooPharma) was injected subcutaneously at 0.5 μg per gram of mouse weight. The dorsal surface was sterilized with iodine and ethanol three times respectively, and four clean, well-defined wounds were created along the middle of the animal’s back using sterile 5 mm biopsy punches (Integra Miltex). Aseptic sticky PDMS silicon ring splints with a 7-mm wide window were adhered to the wounds to prevent skin contraction and allow for wound healing only via re-epithelialization and granulation tissue deposition^57^. The microgels and annealing solution were thoroughly mixed and applied to the wounds immediately. After 30-minute gelation time, animals were carefully wrapped with Tegaderm dressings (3 M, Inc.) and monitored until full recovery from the anesthesia. Animals were housed individually in cages with sufficient enrichments, weighted for the first seven days post-operation, and checked on every other day afterward.

### ELISA

To assess the anti-MMP antibodies, sera collected from mice at designated time points were analyzed for antibody titers by ELISA. Briefly, plates were coated with MMP peptide solution (20 μg /ml) or PBS overnight at 4 C. Plates were washed with PBS containing 0.05% Tween 20 (PBST) and then blocked with PBST containing 2% bovine serum albumin (PBST-BSA) for 1 hour at room temperature. Serum was serially diluted with PBST-BSA in 10-fold steps, applied to coated wells, and incubated for 2 hours at room temperature. To detect total IgG, HRP conjugated Fcγ fragment specific goat anti-mouse IgG (Jackson Immunoresearch) was used as the detection antibody and developed with TMB substrate (Thermofisher). For antibody isotyping, HRP-conjugated IgG subtype-specific, i.e., IgG1, IgG2b, IgG2c, and IgG3, antibodies were utilized (Southern Biotech) in place of the total IgG detection antibody while all other steps were similar. A reported titer of 1 indicates no detectable signal above the background. The optical density at 450 nm was read using a Spectramax i3X microplate reader (Softmax Pro 3.1 software; Molecular Devices).

### Histology staining

At designated time points, the implants or wound samples were extracted for histology examination. For paraffin embedding, samples were fixed with 4% paraformaldehyde overnight at 4 °C before further processing. Paraffin blocks were sectioned into 5 μm thickness with at least 3 serial-sections per slide for hematoxylin and eosin (H&E) staining, masson trichrome staining, Picro-Sirius red staining and immunohistochemistry (IHC) staining.

For H&E staining, paraffin-embedded sections were de-waxed and hydrated using xylene then decreasing ethanol concentrations. They were stained in Mayer Hematoxylin Solution (EMS) for 15 minutes before being rinsed in warm running tap water for 15 minutes. They were placed in DI water for 30 seconds, 95% ethanol for 30 seconds, and then into Alcoholic Eosin Y Counterstain (EMS) for 30 seconds. They were then dehydrated and cleared before being mounted in DPX (EMS).

For Masson’s Trichrome staining, paraffin-embedded sections were de-waxed and hydrated using xylene then decreasing ethanol concentrations. They were mordanted in Bouin’s Fixative (EMS) overnight at room temperature. The slides were rinsed in DI water for 15 minutes. They were then stained in Weigert’s Iron Hematoxylin Working Solution (EMS) for 5 minutes and rinsed in DI water for 10 minutes. The slides were placed in Biebrich Scarlet-Acid Fuchsin (EMS) for 15 minutes and rinsed in DI water for 5 minutes. Next, they were stained in Phosphomolybdic Acid-Phosphotungstic Acid (EMS) for 15 minutes and then in Aniline Blue (EMS) for 10 minutes. The slides were washed in DI water for 5 minutes and differentiated in 1% (v/v) Acetic Acid. They were then dehydrated and cleared before being mounted in DPX (EMS).

For Picro-Sirius red staining, paraffin-embedded sections were de-waxed and hydrated using xylene then decreasing ethanol concentrations. The sections were then stained in Picro-Sirius red (Spectrum Chemical) for one hour. They were then washed two tines in acidified water for 2 minutes each. They were then dehydrated and cleared before being mounted in DPX (EMS)^46^.

For IHC staining, paraffin-embedded slides were deparaffinized with xylene and descendant ethanol. Antigen retrieval was performed by incubating the slides for 20Lminutes in 10 mM sodium citrate buffer with 0.05% Tween 20, pHL6 (VWR) at 95 °C using a microwave. The slides were brought to room temperature and rinsed in PBST (Phosphate Buffered Saline containing 0.05% Tween-20). These slides were stained with rabbit primary antibody (anti-mouse CD11b antibody from Novus Biologicals or anti-mouse α-SMA antibody from Abcam) at 4°C overnight. ImmPRESS® Horse Anti-Rabbit IgG PLUS Polymer Kit (Vector Laboratories) was then used for visualization of CD11b or α-SMA in brown. Subsequently, the slides were washed in tap water, counterstained with Mayer hematoxylin solution (EMS), dehydrated in ethanol, and mounted with DPX (EMS).

### *In vivo* quantification and analysis

For all the stained slides, full section scans were performed using ZEISS Axio Scan.Z1. Images were quantified with an in-house algorithm in ImageJ to determine CD11b+ and α-SMA+ cell area as a ratio of the nucleus area. Briefly, three regions of interest were drawn manually in the connective tissue, inside the remaining biomaterials, and in normal skin, respectively. The total nucleus area was calculated after color deconvolution with H&E DAB vector module and thresholding in the blue channel with ’Max Entropy” method. Another thresholding method (“IsoData”) was implemented in the brown channel to quantify the positive area of CD11b or α-SMA. The expression of each marker was normalized by dividing the positive marker area by the total nucleus area. The remaining MAP scaffolds percentage in the wound was calculated by selecting manually and measuring the area of MAP scaffolds as well as the wound. The epidermis to dermis ratio was measured using an in-house algorithm in ImageJ. Briefly, the dermis area and epidermis area were manually selected using the Freehand Selection tool. The areas were saved and analyzed using a code that measures the width of the epidermis and dermis at multiple matching intervals along the selection. These values were used to create multiple epidermis to dermis ratios and produce a histogram of values with a count, average, and standard deviation. Collagen alignment was measured using were selecting five regions of 100 µm^2^ and the anisotropy was quantified using Fibriltool^59^. Area of collagen, average collagen length and average collagen width were measured on brightfield images of Picro-Sirius red.

These images were thresholding with “IsoData” method, and the masks were used to quantified collagen architecture features.

H&E sections were also examined by a board-certified dermatopathologist (P.O.S.), who was blinded to the identity of the samples, to assess various aspects of wound healing. Re-epithelialization, granulation tissue formation and vascularization, collagen deposition and fibrosis/fibroplasia (early scar formation), and inflammation scores were evaluated by a modified 12-point scoring system^60^ (listed in Tables S1–S5), which was established and agreed upon by two dermatopathologists. Within all MAP scaffolds-treated wounds, we also quantified key markers including the number of hair follicle structures, the number of sebaceous glands, and the number of dermal cysts. Hair follicle-like structures were grouped with utricular pouches, a common structure in SKH-1 mouse skin raised from the infundibulum of the hair shaft (the upper portion of the follicle)^61^. Only the structural features within the inner 60% of the wound were quantified to ensure the regions of interest were within the wound bed and not related to the primary intention of the skin surrounding the injury.

### Statistics and reproducibility

All the statistical analysis was performed using Prism 9 (GraphPad, Inc.) software. Specifically, a one-way or a two-way ANOVA was used to determine the statistical significance. For one-way ANOVA, a post hoc analysis with the Dunnett’s multiple comparisons test was used. For two-way ANOVA, a multiple comparison analysis with the Šídák’s multiple comparisons test was used. The subcutaneous implantation studies (n=5, all-male) were repeated three times and the implants were examined with an 11-color innate flow cytometry panel or a 7-color macrophage panel. The wound healing study (n=5, mix-gender) was repeated twice and the wounds were examined with a 13-color innate flow cytometry panel and/or with histological analysis.

## Supplementary Figures

**Supplementary Figure 1:**
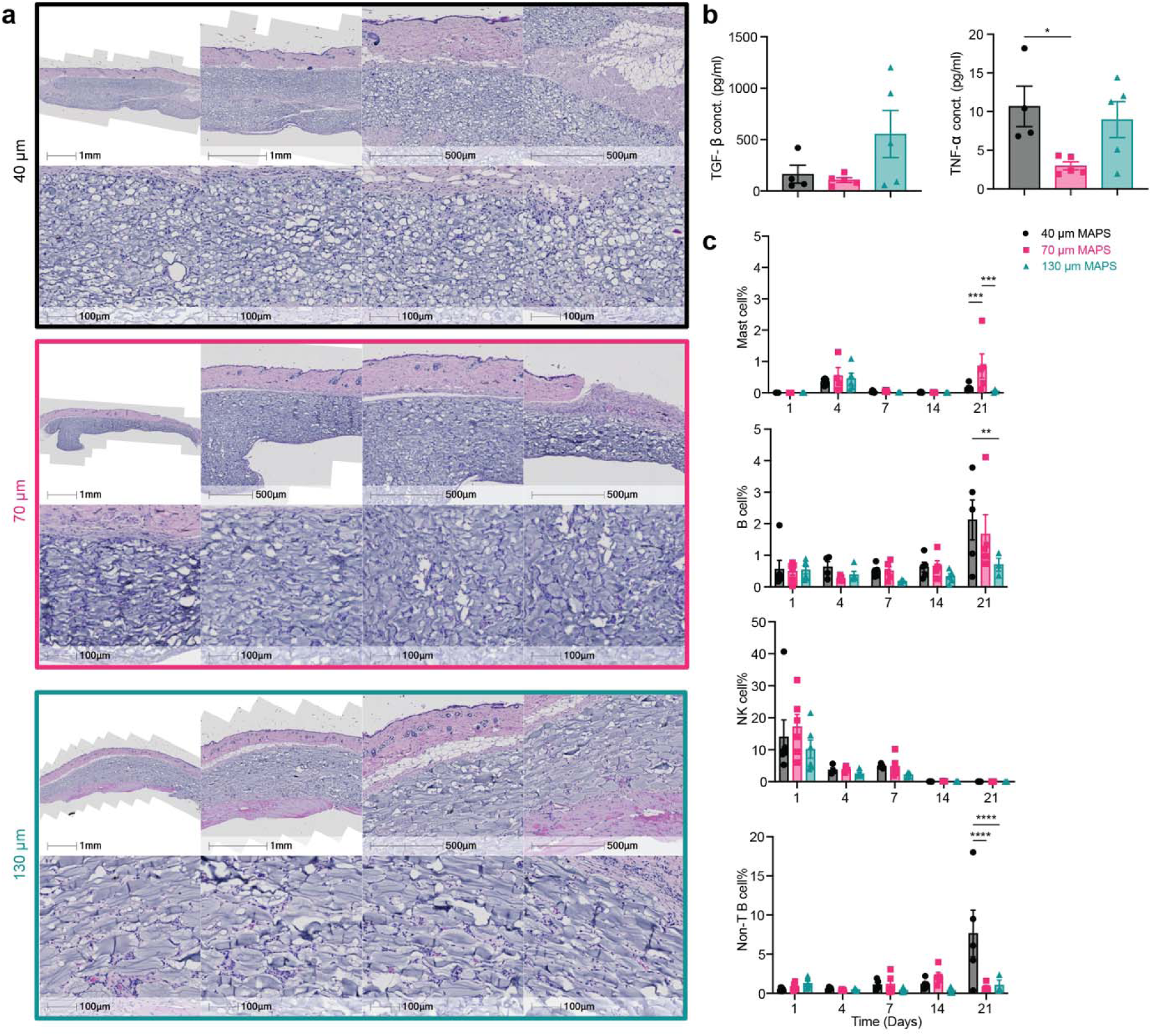
40 μm, 70 μm, and 130 μm MAP scaffolds showed a difference in immune cell recruitment and FBR in the subcutaneous implantation model. a, representative pictures of Hematoxylin and eosin (H&E) staining on day 21. The top row in each panel from left to right showed pictures with objectives of 1x and 2x (implant overview, scale bar, 1 mm), 5x (skin/dorsal interface, scale bar, 500 μm), 5x (capsule/ventral interface, scale bar, 500 μm). The bottom row in each panel showed representative pictures inside the implant with objectives of 10x. b, ELISA results of selected cytokine concentrations inside the hydrogel implants. c, mast cell, B cell, NK cell, non-T/B cell percentages among all CD45+ live cells from flow cytometry. Statistical analysis: two-way ANOVA with Šídák’s multiple comparisons test made between treatment groups only when there was a significance in the interaction term of treatment type x time. ∗ p < 0.05, ∗∗ p < 0.001, ∗∗∗ p < 0.001, ∗∗∗∗ p < 0.0001. Error bars, mean ± s.e.m., n = 5 mice per group with some data points removed due to experimental reasons.

**Supplementary Figure 2:**
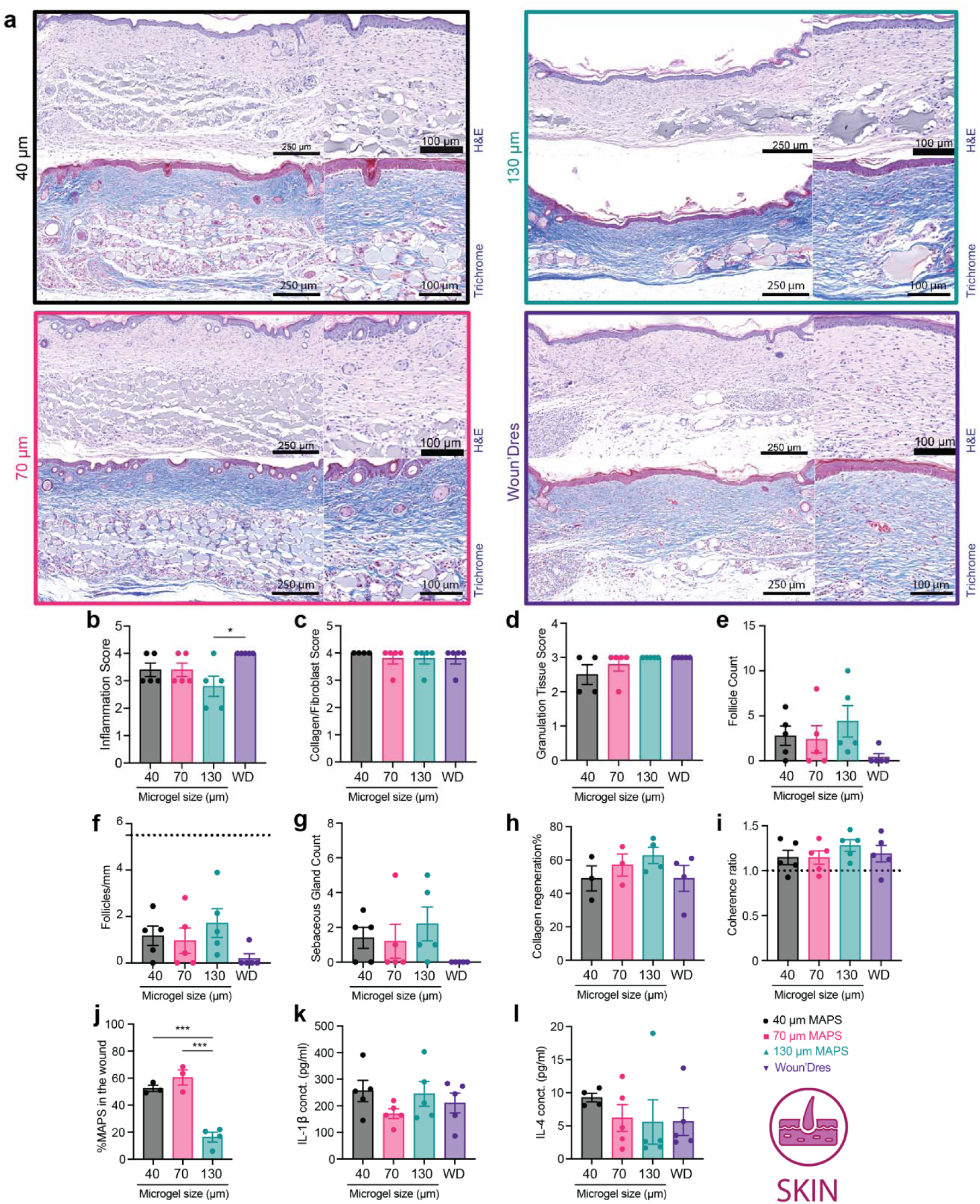
130 μm MAP scaffolds induced mature collagen regeneration and reduced inflammation level in the skin wound. a, representative pictures of 21-day skin wound samples with H&E staining and Masson’s trichrome staining (left panel, full wound and surrounding skin, scale bar, 250 μm; right panel, zoom-in picture at the wound site, scale bar, 100 μm). b-d, histologic assessment of inflammation, collagen/fibroblast score and granulation tissue. e, the cell nucleus area in the connective tissue. The dotted line and the grey area stand for the average number and the range for normal skin. f, the cell nucleus area in the remaining biomaterials. The dotted line and the grey area stand for the average number and the range for normal skin. g-i, histologic quantification of follicle count, follicles/mm, and sebaceous gland count. The gray line stands for the value in normal skin. j, collagen regeneration percentage compared to the normal skin. k, the coherence ratio in collagen fibers. The dotted line stands for the level of normal skin. l, the remaining MAP scaffolds amount in the wound relative to the size of the wound. m-n, ELISA results of IL1-β and IL-4 concentrations inside the hydrogel implants. Statistical analysis: two-way ANOVA with Šídák’s multiple comparisons test made between treatment groups only when there was a significance in the interaction term of treatment type x time. Dunnet method was used to compare each treatment against normal skin baseline (gray pond). ∗/# p < 0.05, ∗∗ p < 0.001, ∗∗∗ p < 0.001, ∗∗∗∗ p < 0.0001. Error bars, mean ± s.e.m., n = 5 mice per group with some data points removed due to experimental reasons.

**Supplementary Figure 3.**
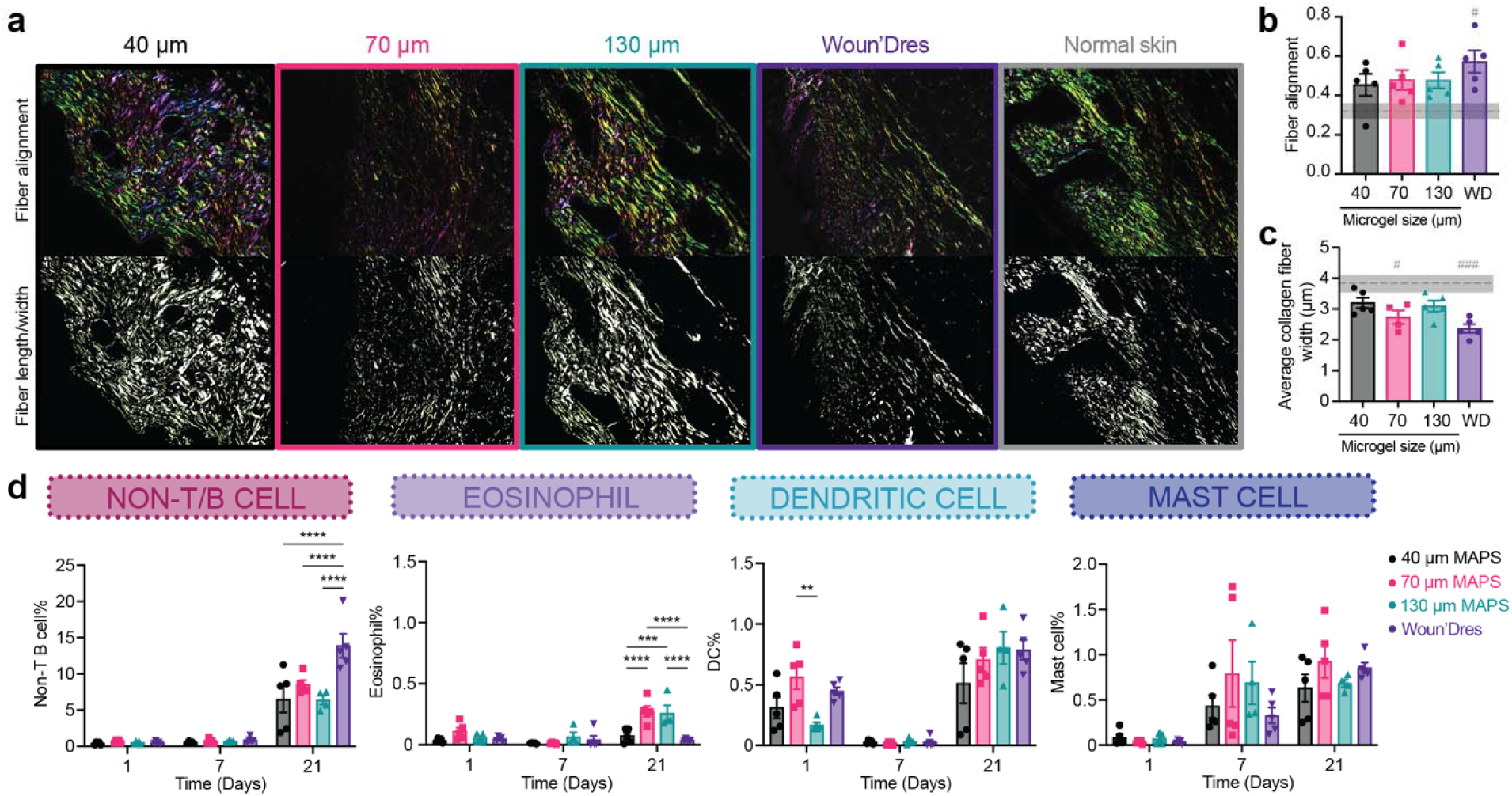
130 μm MAP scaffolds induced mature collagen regeneration, and immune cell recruitment followed a size-dependent manner in the skin wound. a, representative pictures of 21-day skin wound samples with Picro-Sirius Red staining (top row, fiber alignment analysis; bottom row, fiber length/width analysis). b, fiber alignment score (calculated by ImageJ software) in the regions of interest. The dotted line and the grey area stand for the average number and the range for normal skin. c, average width of collagen fibers in the regions of interest. The dotted line and the grey area stand for the average number and the range for normal skin. d, non-T/B cell, eosinophil, dendritic cell, mast cell percentages among all CD45+ live cells from flow cytometry. Statistical analysis: two-way ANOVA with Šídák’s multiple comparisons test made between 40 μm, 70 μm, and 130 μm MAP scaffolds and wound dressing groups only when there was a significance in the interaction term of scaffold type x time. ** p<0.01, *** p<0.001, **** p<0.0001. Error bars, mean ± s.e.m., n = 5 mice per group.

**Supplementary Figure 4:**
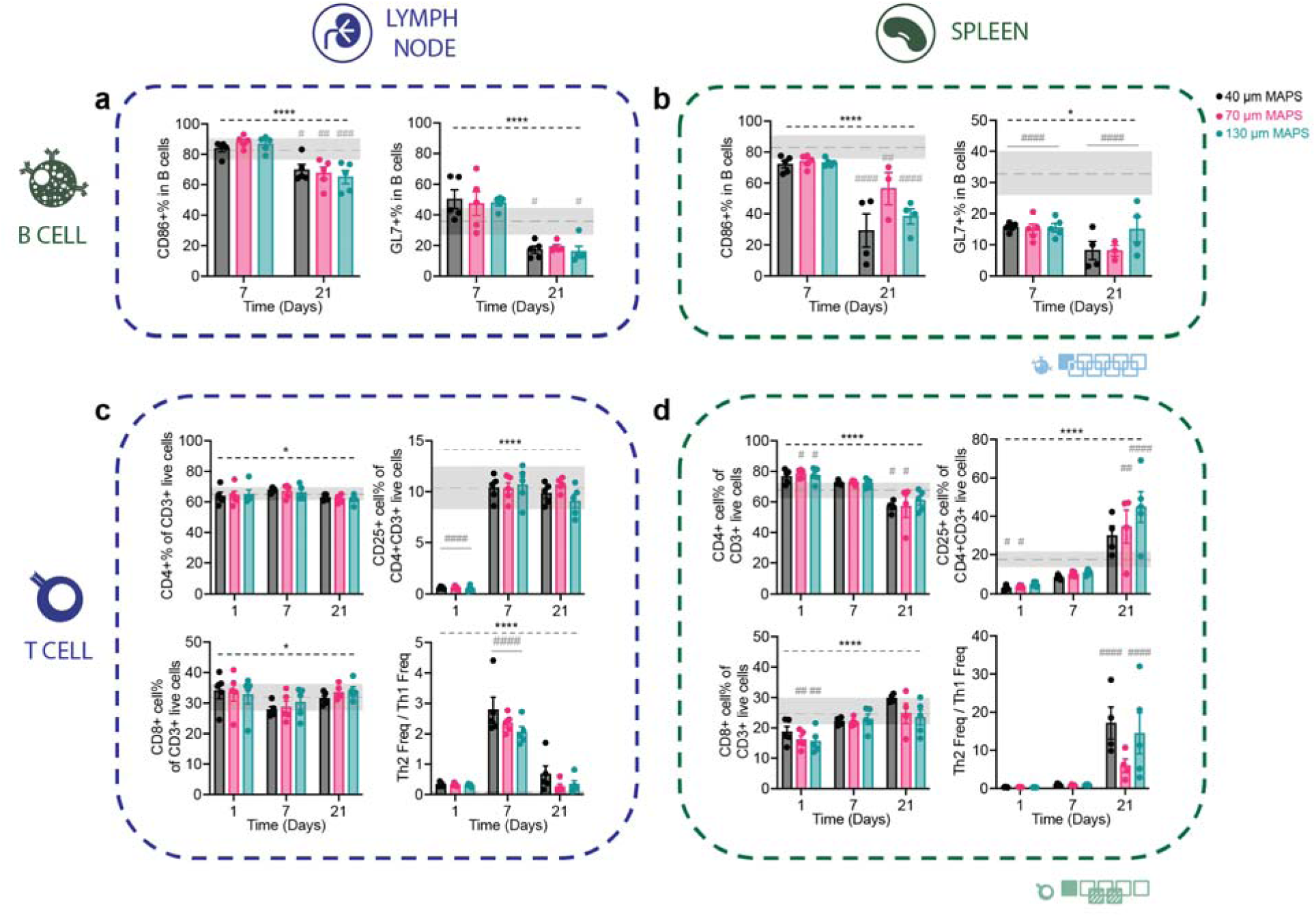
MAP scaffolds treatment induced phenotype switches in both B cell and T cell populations compared to baseline in draining lymph nodes and spleen of mice. a and b, B cell profiles in draining lymph nodes and spleen of mice treated with 40 μm, 70 μm, and 130 μm MAP scaffolds across 3 time points. c and d, T cell profiles in draining lymph nodes and spleen of mice treated with 40 μm, 70 μm, and 130 μm MAP scaffolds across 3 time points. Statistical analysis: two-way ANOVA with Šídák’s multiple comparisons test made between 40 μm, 70 μm, and 130 μm MAP scaffolds groups only when there was a significance in the interaction term of scaffold type x time. After a two-way ANOVA, Dunnet method was used to compare the experiment groups with the baseline control group (mice without wounding). */# p<0.05, **/## p<0.01, ***/### p<0.001, ****/#### p<0.0001. Asterisks with solid line stand for comparisons between MAP scaffolds. Asterisks with dash line stand for significant difference in time. Error bars, mean ± s.e.m., n = 5 mice per group but with some data points removed due to experimental reasons. The blue symbol beneath panel b stands for the 9-color B panel used in panel a, b of this figure. The green symbol beneath panel d stands for the 7-color T cell panel used in panel c, d of this figure.

**Supplementary Figure 5:**
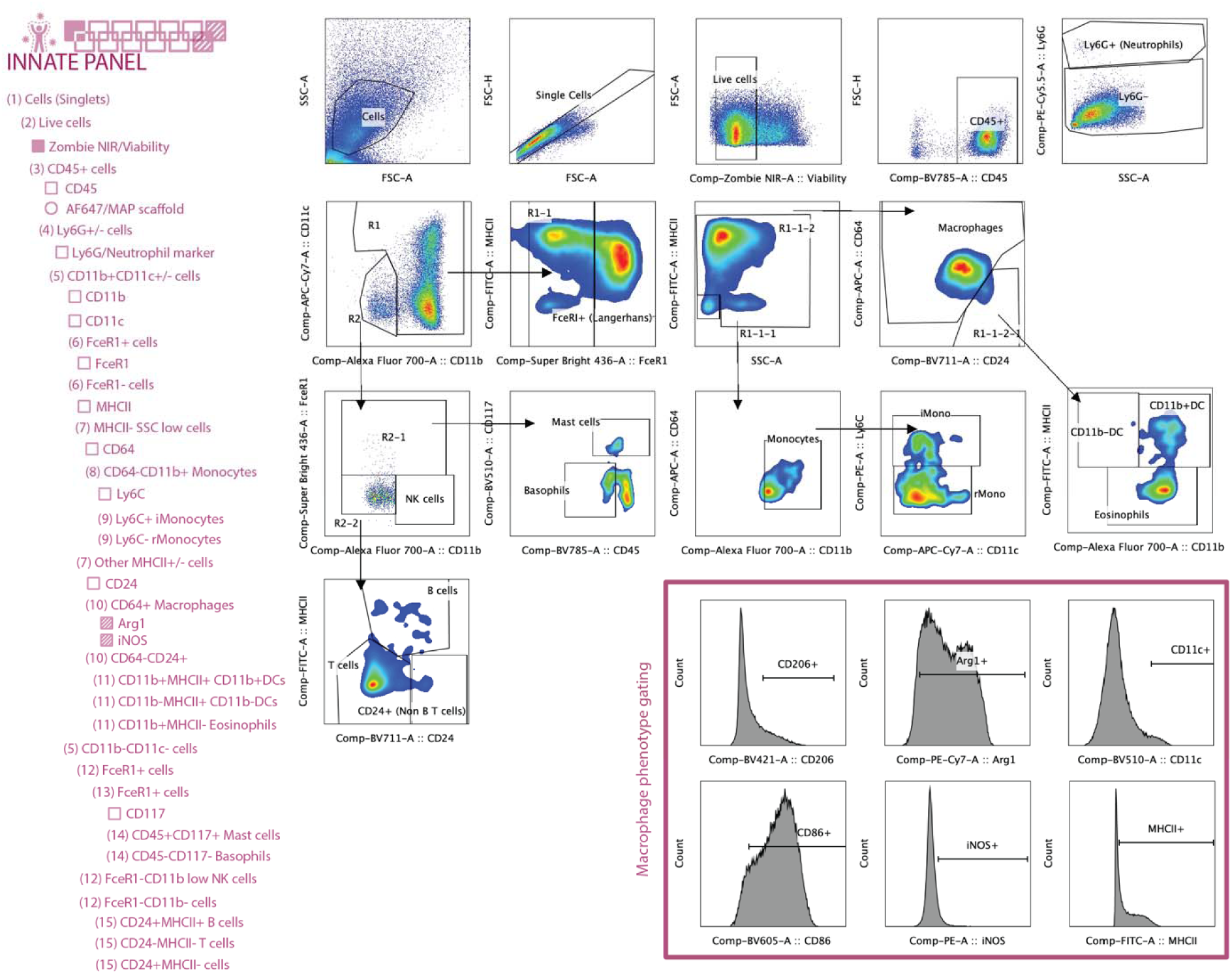
Gating strategy for flow cytometry analysis on myeloid cells with 13 markers.

**Supplementary Figure 6:**
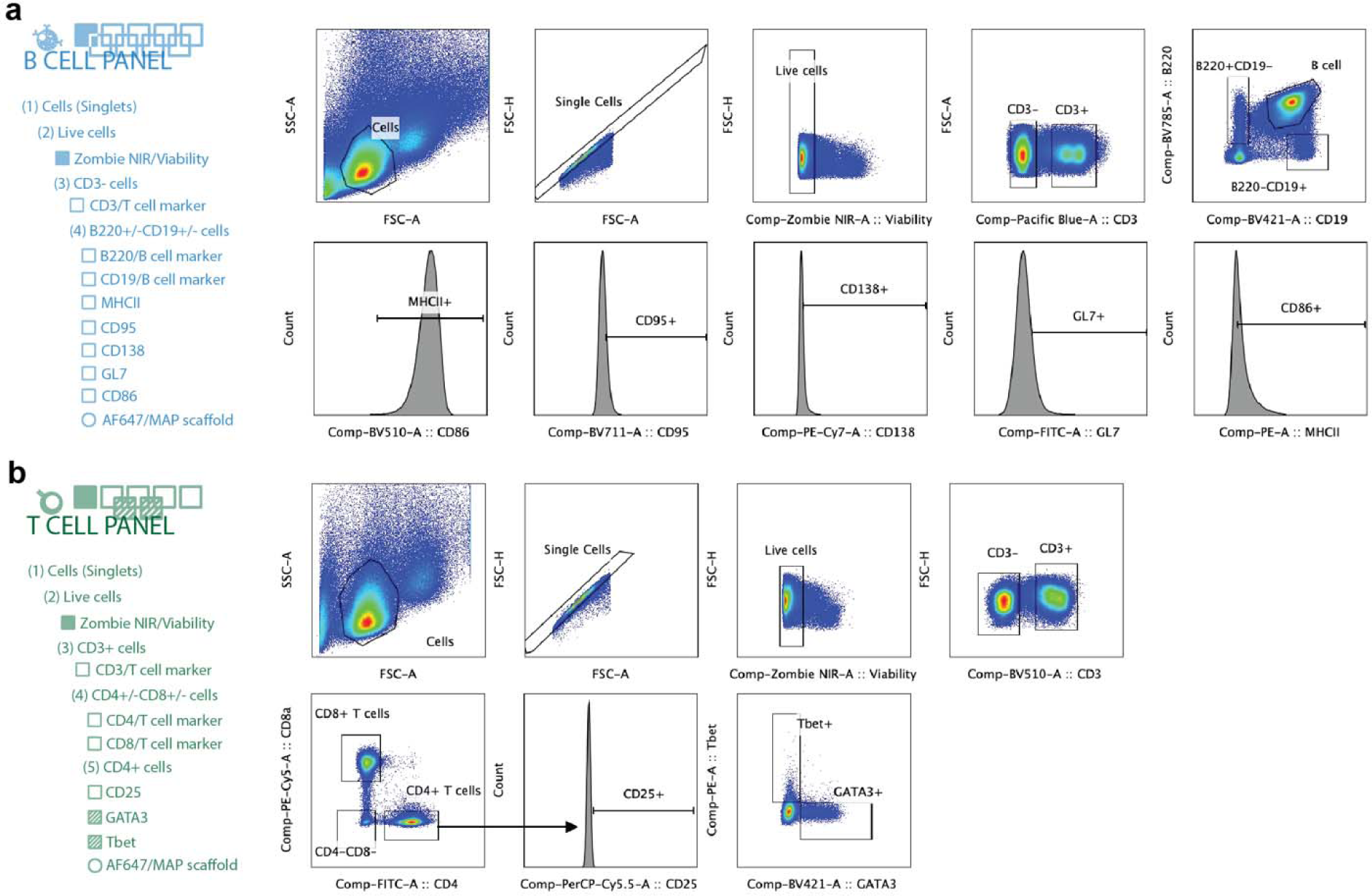
Gating strategy for flow cytometry analysis on: a) B cells with 9 markers. b) T cells with 7 markers.

**Supplementary Table 1.**
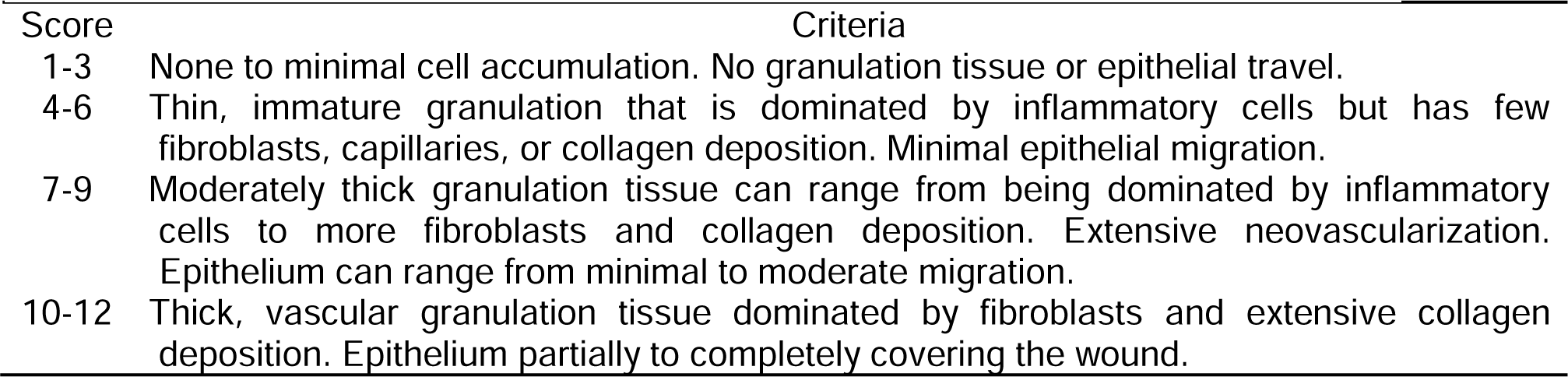
Scoring of Histological Sections for Total Wound Healing

**Supplementary Table 2.**
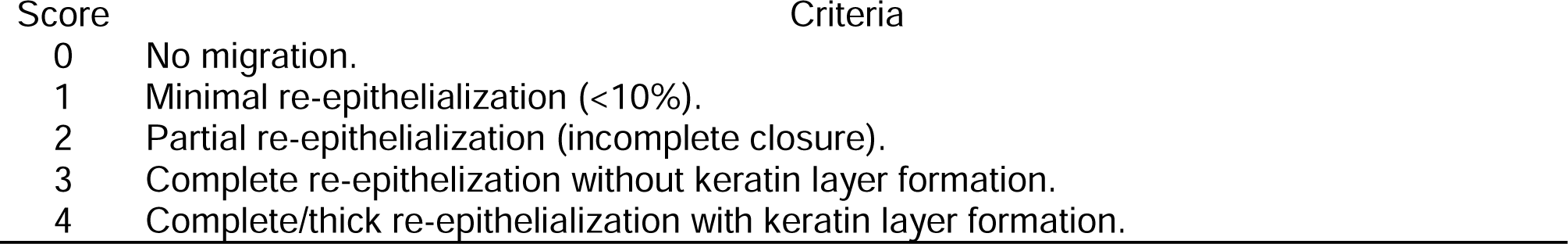
Scoring of Epidermis/re-epithelialization

**Supplementary Table 3.**
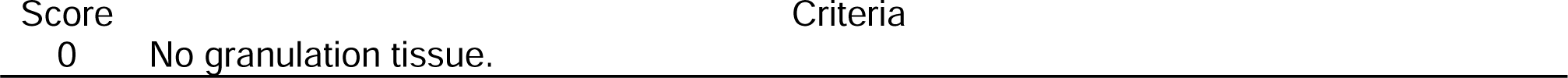
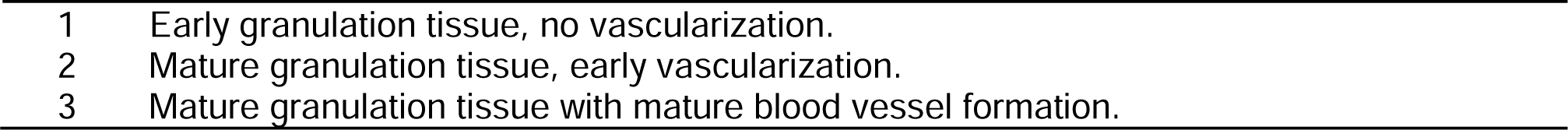
Scoring of Granulation Tissue/Vascularization

**Supplementary Table 4.** Scoring of Collagen Deposition/Fibroplasia

| Score | Criteria |
| --- | --- |
| 0 | No collagen deposition/fibroplasia. |
| 1 | Fibroblast proliferation/no collagen deposition. |
| 2 | Fibroblast proliferation with minimal collagen deposition. |
| 3 | Fibroblast proliferation with extensive haphazard collagen deposition. |
| 4 | Extensive organized collagen deposition or complete replacement of dermis with fibrous tissue (mature scar). |

**Supplementary Table 5.**
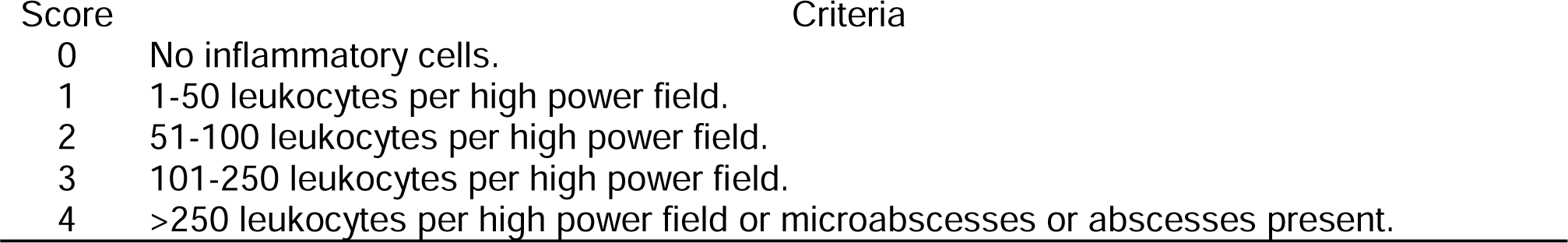
Scoring of Inflammation

**Supplementary Table 6.**
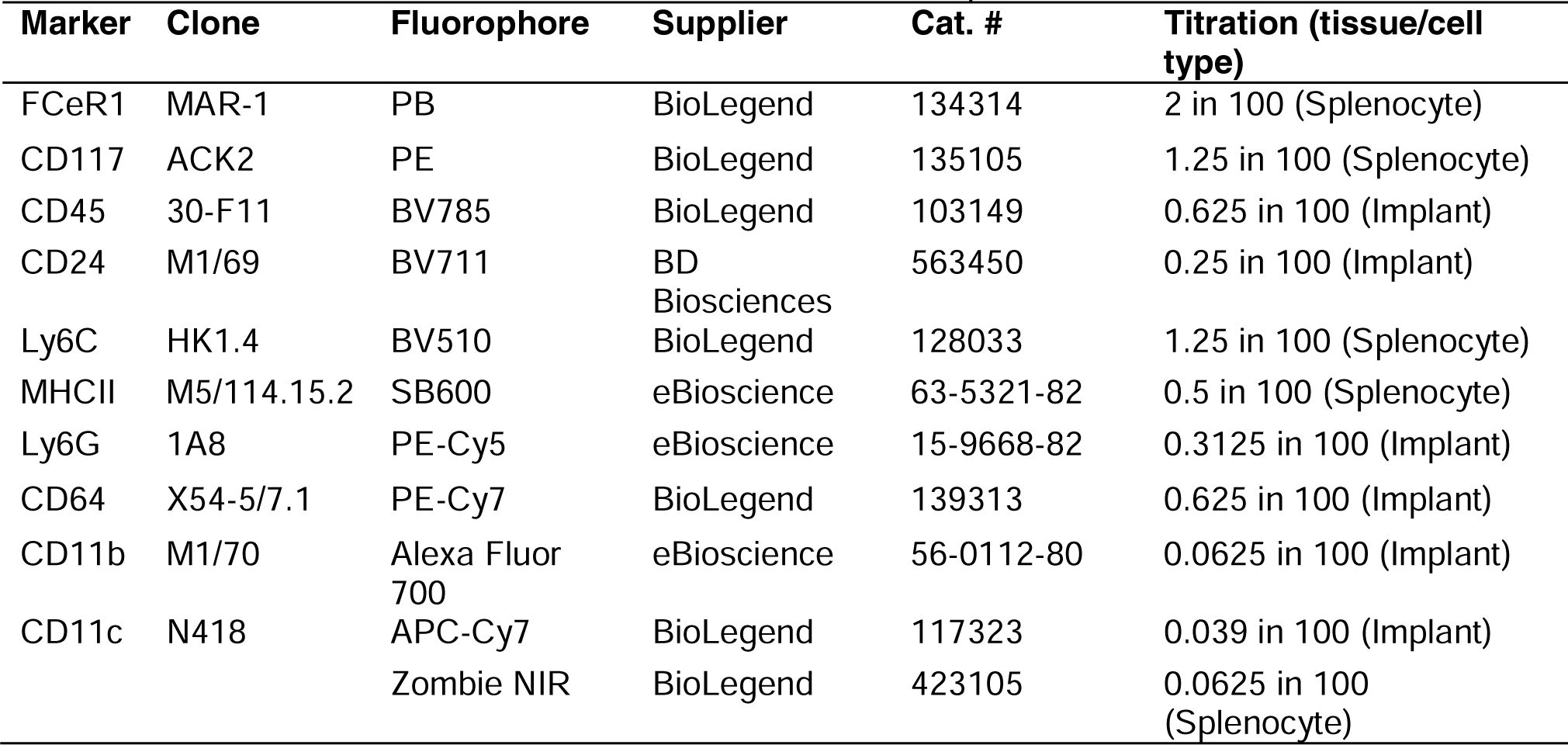
A list of antibodies for the innate panel

**Supplementary Table 7.**
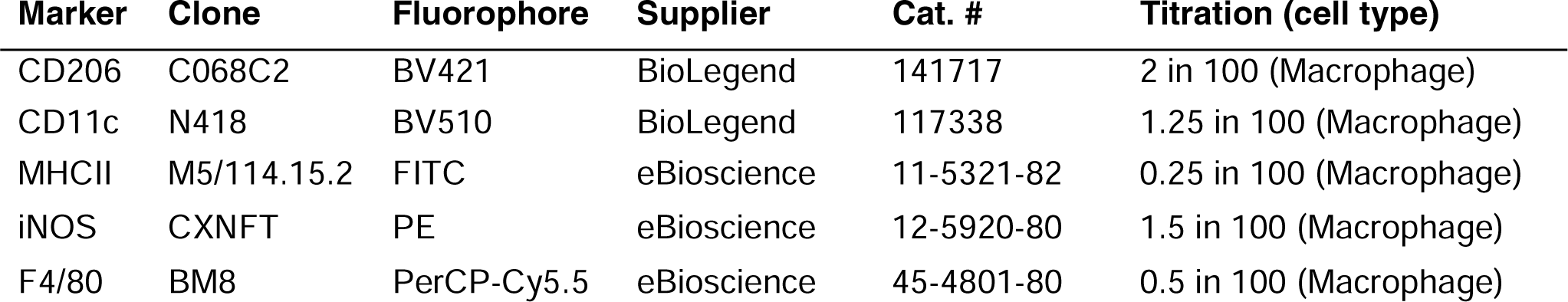
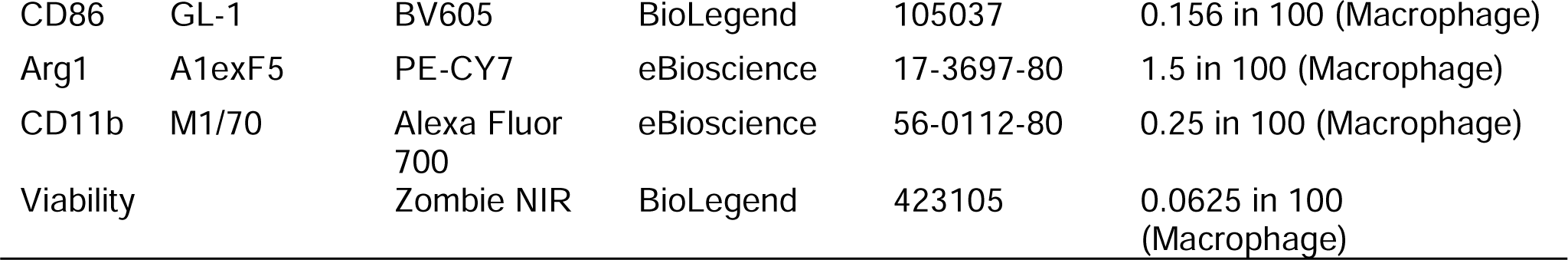
A list of antibodies for the macrophage panel

**Supplementary Table 8.**
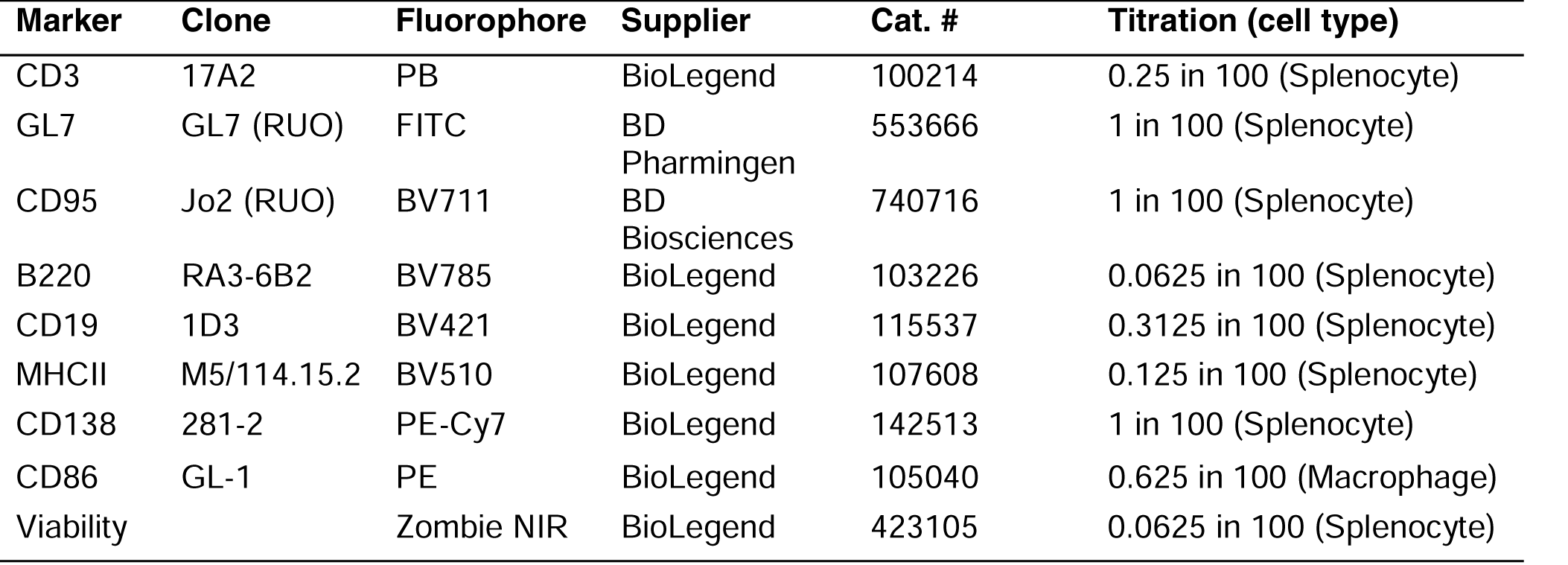
A list of antibodies for the B cell panel

**Supplementary Table 9.** A list of antibodies for the T cell panel

| Marker | Clone | Fluorophore | Supplier | Cat. # | Titration (cell type) |
| --- | --- | --- | --- | --- | --- |
| CD3 | 17A2 | BV510 | BioLegend | 100233 | 1.25 in 100 (Splenocyte) |
| CD4 | GK1.5 | FITC | BD Pharmingen | 557307 | 0.156 in 100 (Splenocyte) |
| CD8a | 53-6-7 | PE-Cy5.5 | BioLegend | 100710 | 0.1 in 100 (Splenocyte) |
| CD25 | PC61 | PerCP-Cy5.5 | BioLegend | 102030 | 0.5 in 100 (Splenocyte) |
| Tbet | 4B10 | PE | BioLegend | 644810 | 2.5 in 100 (Splenocyte) |
| GATA3 | 16E10A23 | BV421 | BioLegend | 653806 | 2.5 in 100 (Splenocyte) |
| Viability |  | Zombie NIR | BioLegend | 423105 | 0.0625 in 100 (Splenocyte) |

**Supplementary Table 10.** A list of antibodies for the APC panel

| Marker | Clone | Fluorophore | Supplier | Cat. # | Titration (cell type) |
| --- | --- | --- | --- | --- | --- |
| CD3 | 17A2 | NovaFluor Red 685 | Thermo Fisher | M002T02R02 | 0.0625 in 100 (Splenocyte) |
| CD45 | 30-F11 | Alexa Fluor 532 | Thermo Fisher | 58-0451-82 | 0.3125 in 100 (Splenocyte) |
| XCR1 | ZET | APC-Cy7 | BioLegend | 148223 | 0.3125 in 100 (Splenocyte) |
| CD64 | X54-5/7.1 | BV711 | BioLegend | 139311 | 0.156 in 100 (Splenocyte) |
| Ly6C | HK1.4 | BV570 | BioLegend | 128030 | 0.625 in 100 (Splenocyte) |
| CD169 | 3D6.112 | PE-Cy7 | BioLegend | 142412 | 2.5 in 100 (Splenocyte) |
| F4/80 | BM8 | PB | Thermo Fisher | MF48028 | 2 in 100 (Splenocyte) |
| CD80 | 16-10A1 | PE-CF594 | BioLegend | 104737 | 1.25 in 100 (Splenocyte) |
| CD103 | M290 | VioBright515 | Miltenyi | 130-111-609 | 2 in 100 (Splenocyte) |
| CD24 | M1/69 | SB600 | Thermo Fisher | 63-0242-80 | 0.625 in 100 (Splenocyte) |
| CD11c | N418 | SB780 | Thermo Fisher | 78-0114-82 | 0.625 in 100 (Splenocyte) |
| pDCA-1 | 927 | BV650 | BioLegend | 127019 | 0.078 in 100 (Splenocyte) |
| ICAM-1 | 3E2 | SB436 | Thermo Fisher | 62-0542-80 | 0.625 in 100 (Splenocyte) |
| CCR7 | 4B12 | PE | Thermo Fisher | 12-1971-80 | 1.25 in 100 (Splenocyte) |
| B220 | RA3-6B2 | APC/Fire810 | BioLegend | 103277 | 0.3125 in 100<br>(Splenocyte) |
| CD172a | P84 | BB700 | BD Bioscience | 742205 | 1 in 100 (Splenocyte) |
| MHCII | M5/114 | BV510 | BioLegend | 107635 | 0.25 in 100 (Splenocyte) |
| Ly6G | 1A8 | BV480 | BD Bioscience | 746448 | 0.125 in 100 (Splenocyte) |
| CD11b | M1/70 | Alexa Fluor 700 | eBioscience | 56-0112-80 | 0.0625 in 100<br>(Splenocyte) |
| Viability |  | Zombie NIR | BioLegend | 423105 | 0.03125 in 100<br>(Splenocyte) |

**Supplementary Table 11.**
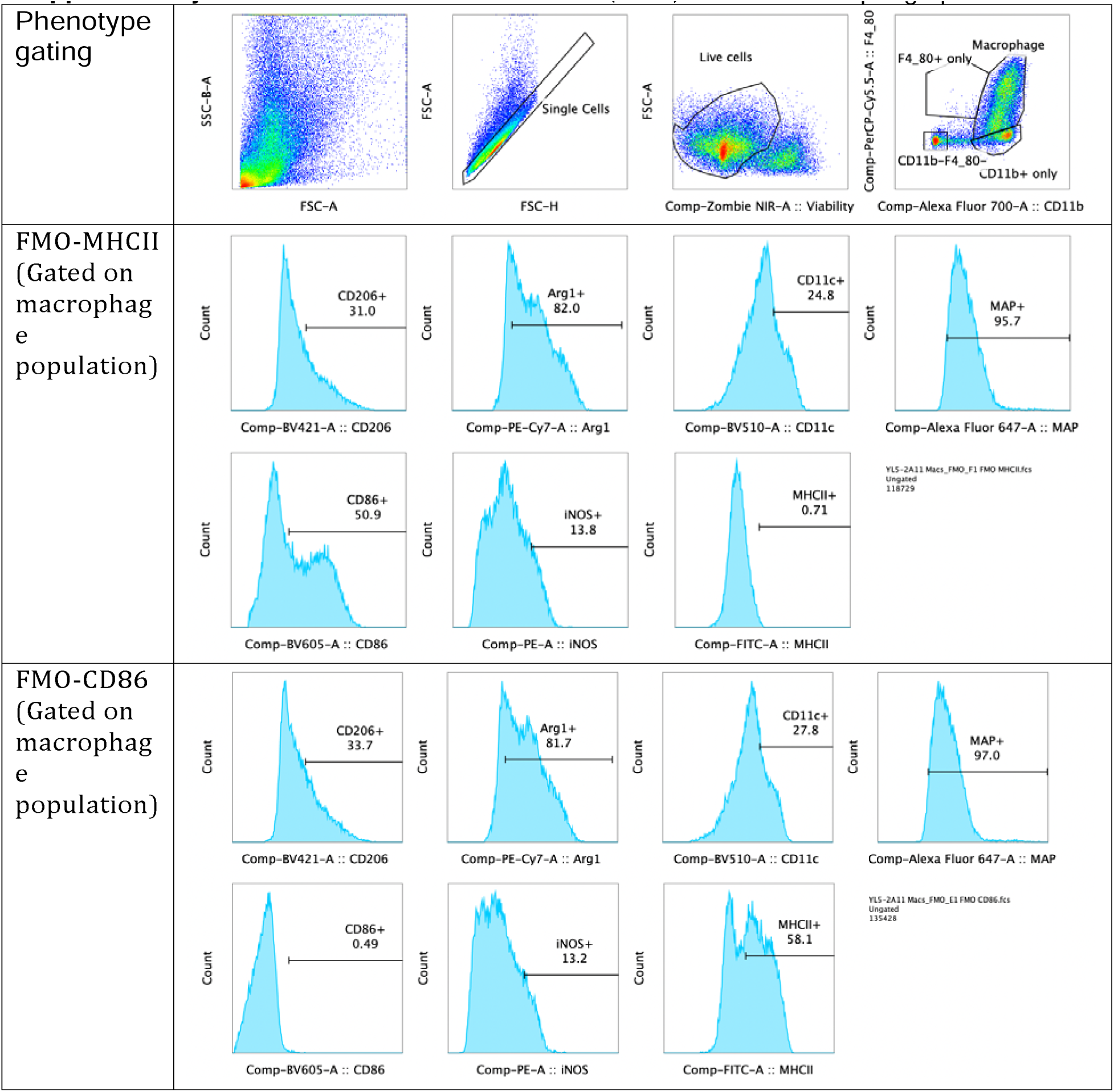
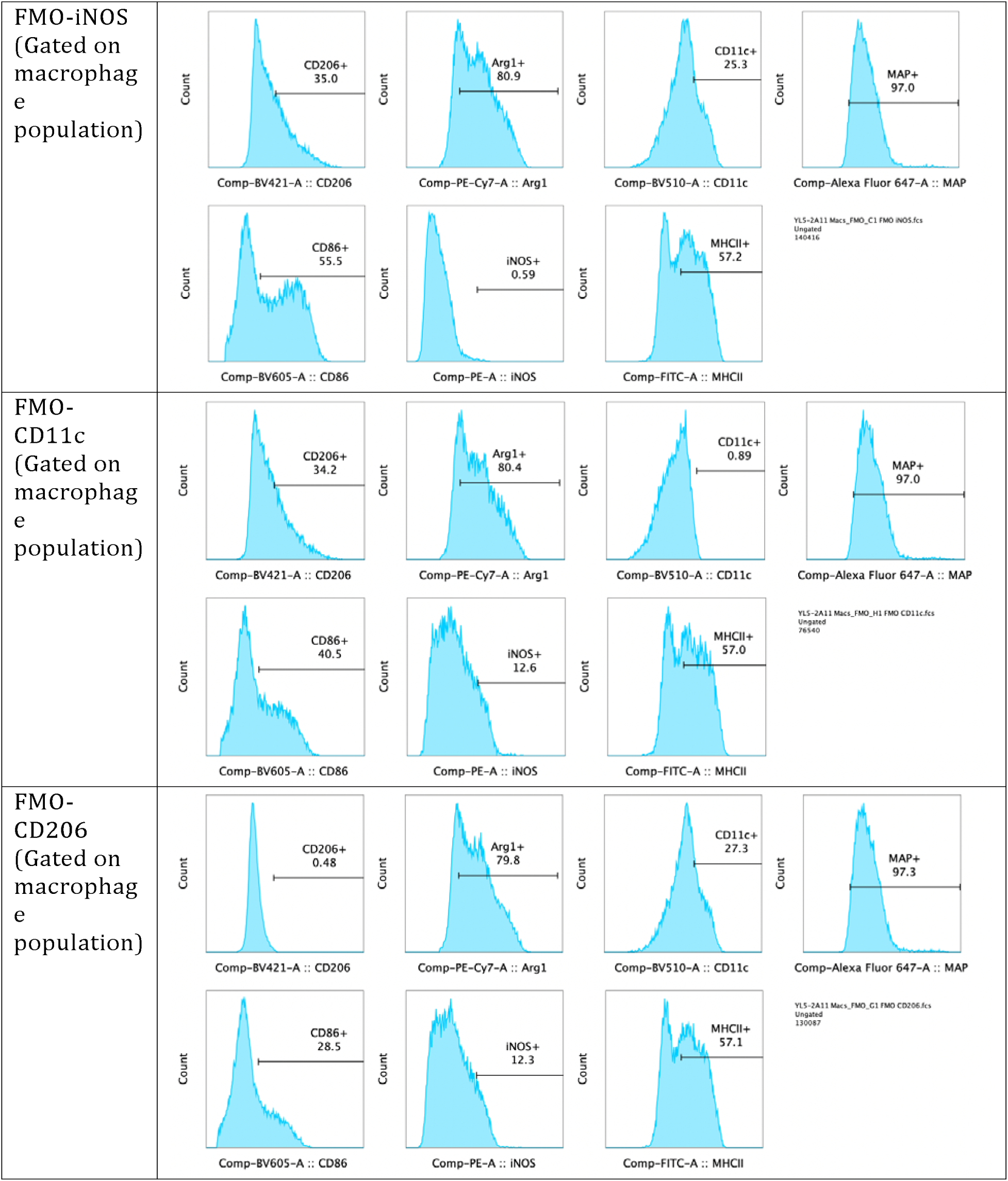
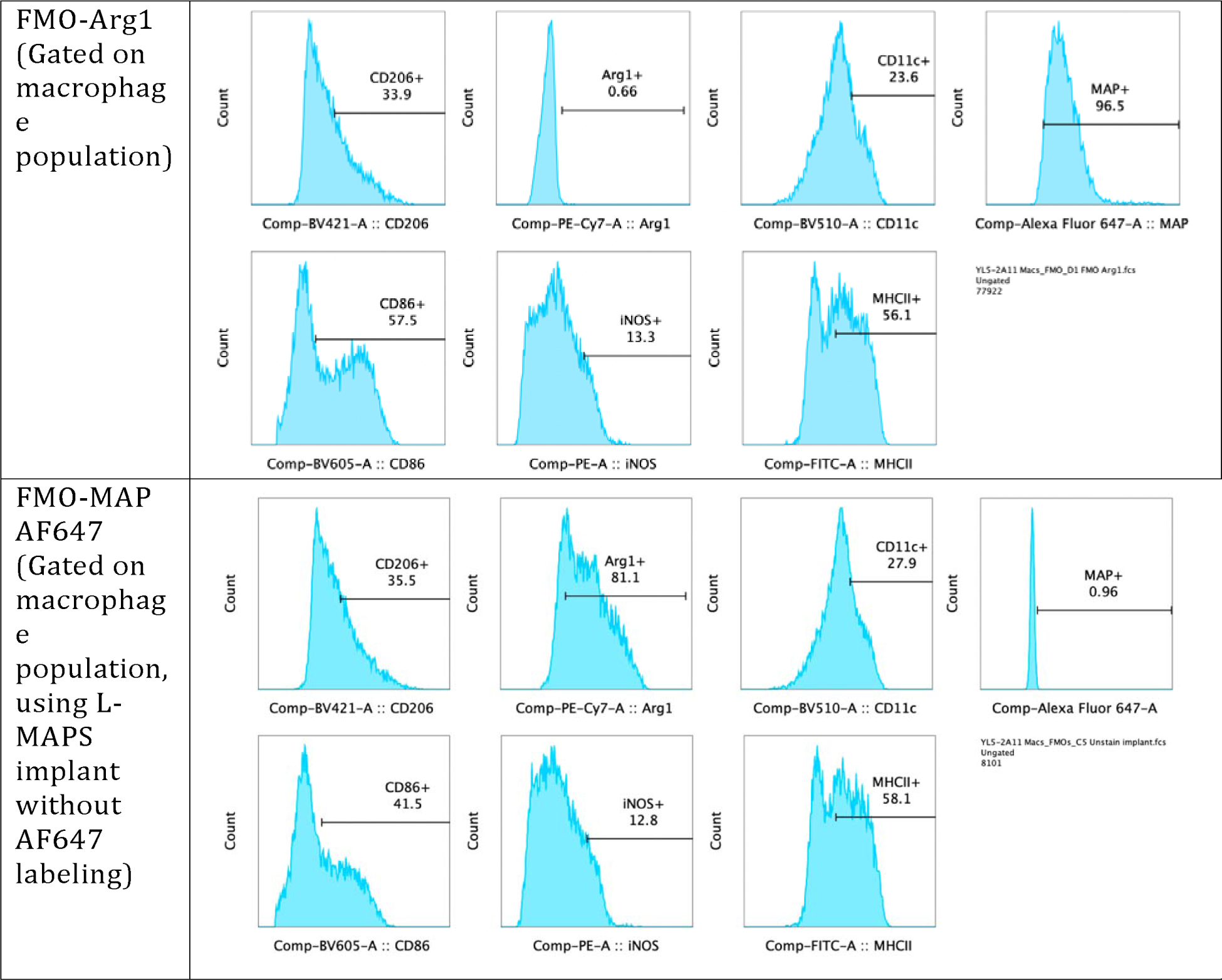
fluorescence minus one (FMO) controls macrophage panel

**Supplementary Table 12.**
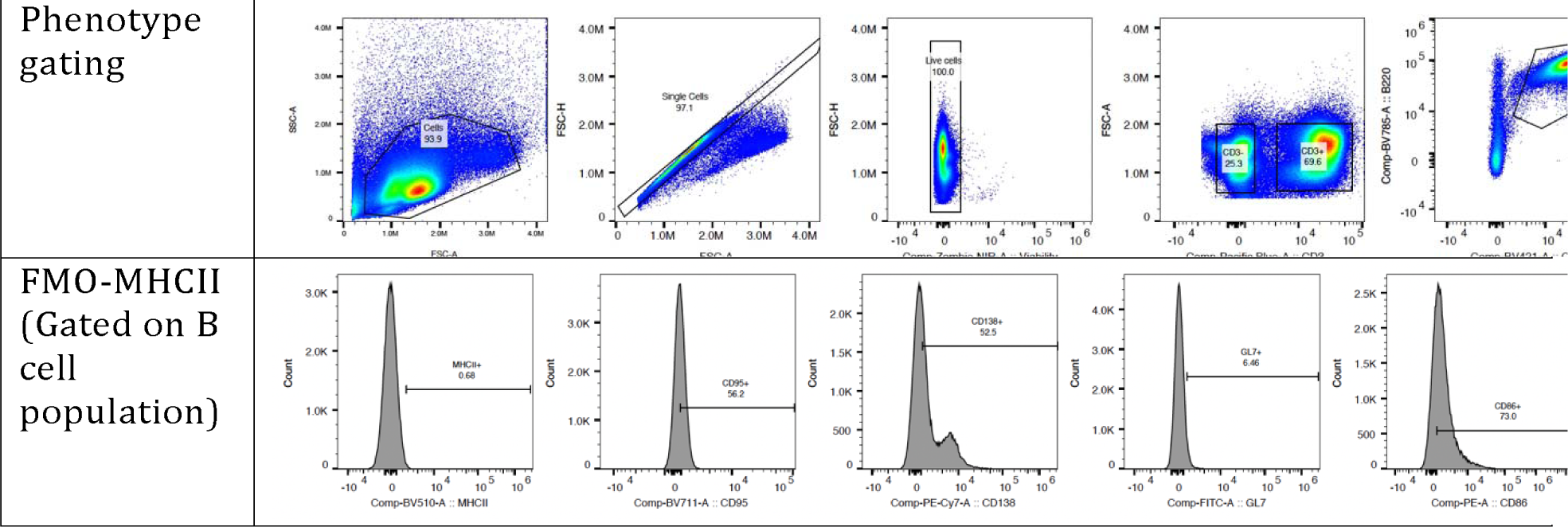
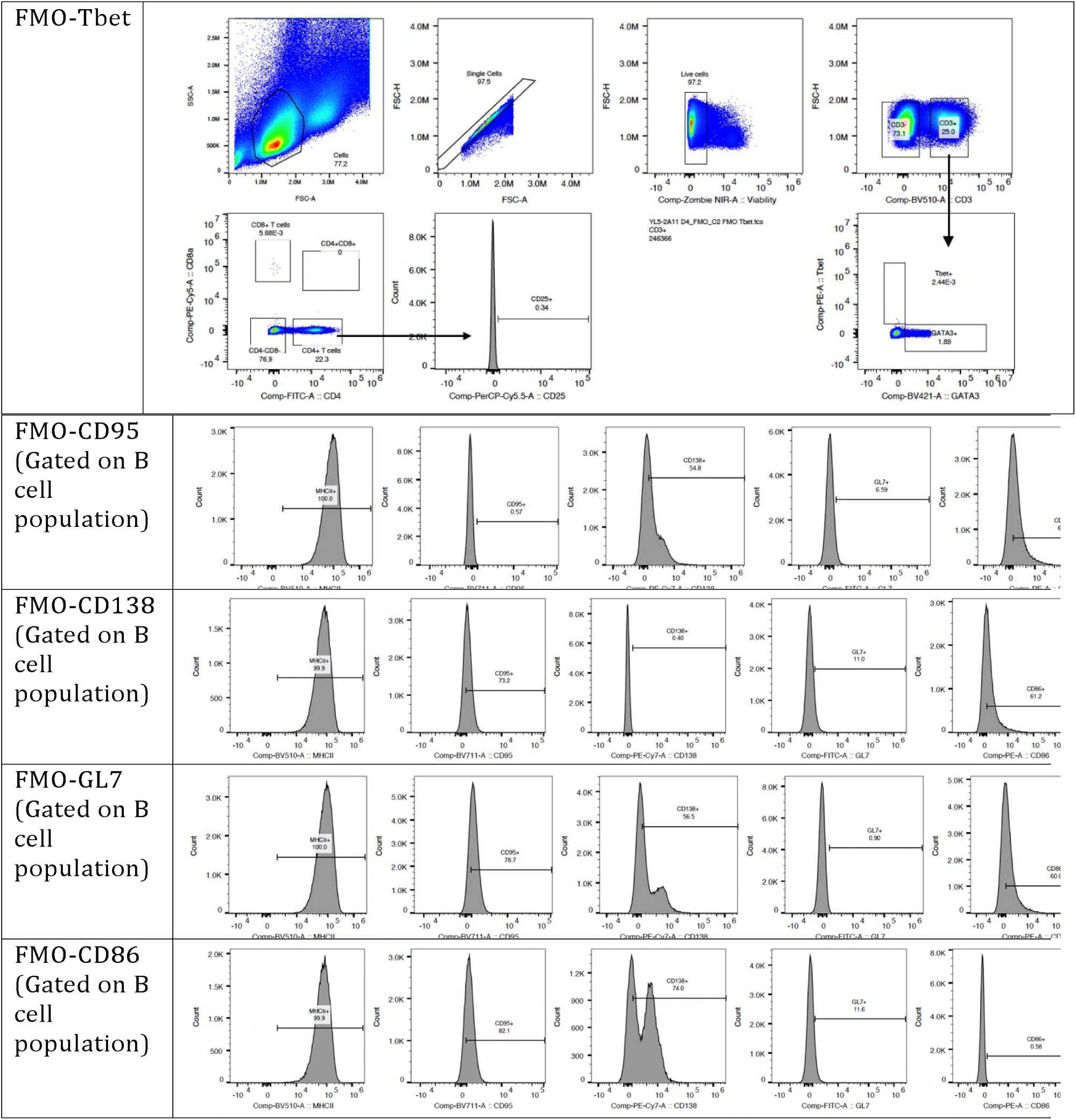
fluorescence minus one (FMO) controls B cell panel

**Supplementary Table 13.**
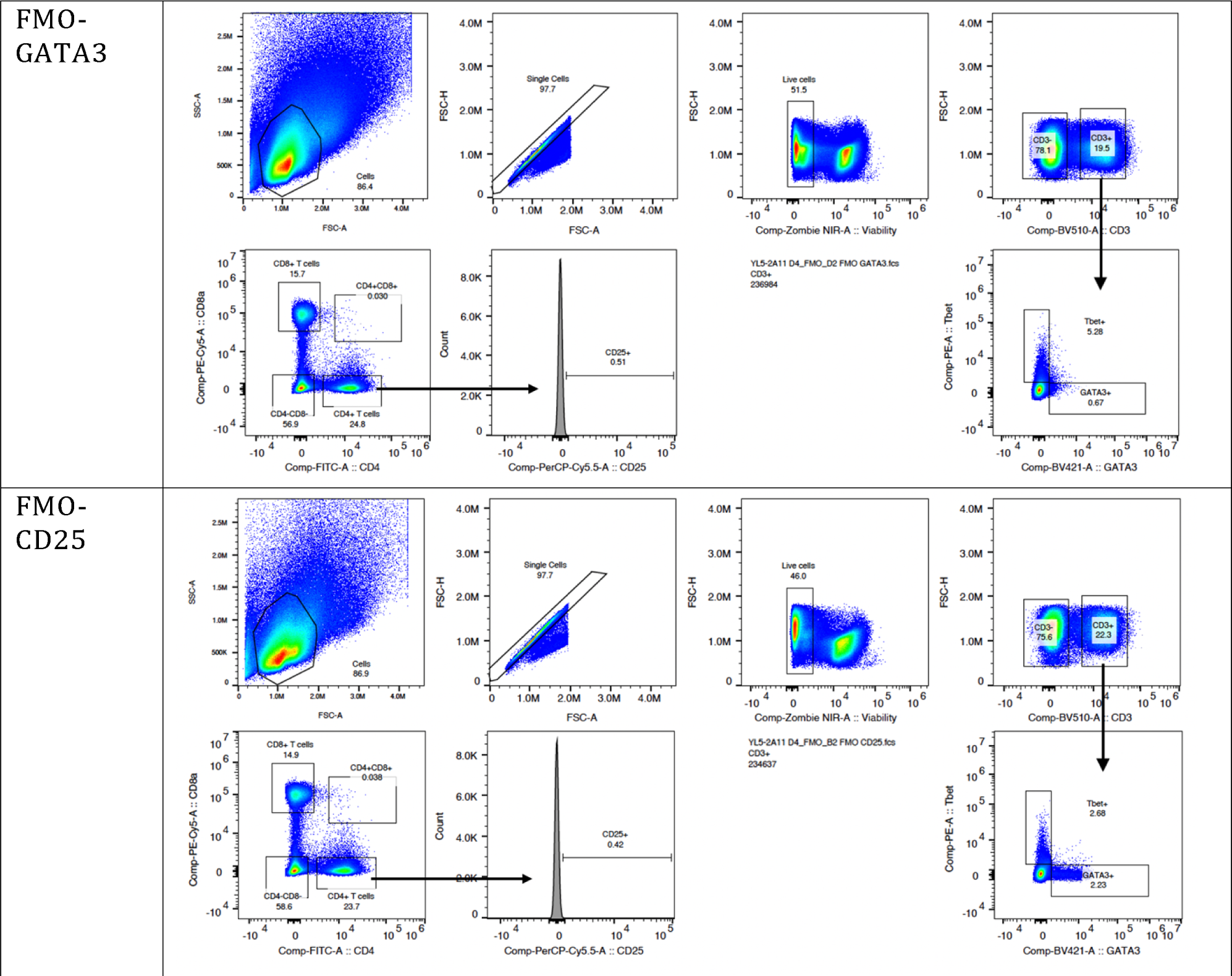
fluorescence minus one (FMO) controls T cell panel

